# Modular in vivo engineering of the reductive methylaspartate cycles for synthetic CO_2_ fixation

**DOI:** 10.1101/2025.11.12.688011

**Authors:** Vittorio Rainaldi, Helena Schulz-Mirbach, Joseph B. Solomon, Stephanie Q. Sloothaak, Nicole Paczia, Tobias J. Erb, Sarah D’Adamo, Nico J. Claassens

**Affiliations:** Laboratory of Microbiology, Wageningen University, Stippeneng 4, 6708 WE, Wageningen, the Netherlands; Bioprocess Engineering, Wageningen University, Droevendaalsesteeg 1, 6708 PB, Wageningen, the Netherlands; Max Planck Institute for Terrestrial Microbiology, Karl-von-Frisch-Str. 10, 35043 Marburg, Germany

## Abstract

Biological carbon fixation is currently limited to seven naturally occurring pathways. Synthetic carbon fixation pathways have the potential to surpass aerobic natural pathways in efficiency, but none have been realized in living cells. Here, we present the reductive methylaspartate cycles (rMASP), a novel family of 4 energy-efficient aerobic CO_2_ fixation pathway variants. These cycles have the potential to outperform the yields and/or rates of the Calvin cycle and previously proposed synthetic CO_2_ fixation cycles. To realize these designs, we adopted a modular engineering approach in *Escherichia coli*. Several pathway modules up to a cascade of 11 enzymes were realized via engineering and evolution. We demonstrate crotonyl-CoA carboxylase dependent growth for both elevated and ambient CO_2_ conditions, and show in vitro activity of the other CO_2_-fixing 2-oxoglutarate carboxylase, which requires further activity optimization for mesophilic temperatures. This work demonstrates important steps toward realizing efficient synthetic carbon fixation pathways in living organisms.

---

Biological carbon fixation is crucial for life on earth, assimilating 380 billion tons of CO_2_ per year^1^. Seven CO_2_ fixation pathways have been discovered so far in nature, all with different architectures, physiological properties, and ecological niches^2,3^. Of these seven, four require oxygen-sensitive enzymes and are thus restricted to anaerobic environments^4^. Under aerobic conditions, the Calvin-Benson-Bassham (CBB) cycle is by far the most common and is responsible for the vast majority of biological carbon fixation on earth^5^.

Despite its prevalence the CBB cycle suffers from several issues that decrease its efficiency, activity, and overall yield. The key enzyme of the CBB cycle, ribulose bisphosphate carboxylase/oxygenase (rubisco), is known to be kinetically slow^6–9^, and as a result it must be produced in large quantities, making rubisco the most abundant enzyme on earth^10^. Furthermore, rubisco shows a seemingly unavoidable side-activity with oxygen, requiring expensive metabolic strategies to deal with the toxic byproduct of the oxygenation reaction, a process known as photorespiration^11^, which results in a yield loss of 20-36% in C3 crops such as soy^12^. While countless efforts have been made to improve rubisco, most of them have failed or ran into the trade-off of improving one feature (e.g. kinetic rate) and worsening another (e.g. oxygenase side-activity)^13^.

To bypass rubisco, researchers have proposed numerous alternative pathway architectures for carbon fixation by combining known biochemical reactions in ways that have not been discovered in nature^14–17^. According to theoretical calculations, some of these may lead to more efficient carbon fixation, by saving ATP-costs and/or eliminating photorespiration losses^16^. These pathways focus on carboxylases other than rubisco, with the intent of eliminating the kinetic bottleneck and the oxygen side-activity that are associated with it. While dozens of different pathways have been proposed, most have never been validated experimentally, few have been tested in vitro^18–20^ and only one has been partially engineered in vivo^21^. In contrast, routes for the assimilation of more reduced C1 compounds such as formate^22,23^ or methanol^24,25^ were successfully realized in vivo, allowing for full biomass formation and even production from these substrates^26,27^. While these results have potential for microbial C1 valorization, feeding reduced substrates is not applicable to plants and algae, which natively capture CO_2_ while harnessing light energy. Realizing the proposed metabolic routes for the direct and efficient use of atmospheric CO_2_ could overcome this limitation. To date, the CBB cycle is the only full oxygen-tolerant CO_2_ fixation pathway that has been successfully introduced in heterologous hosts such as *Escherichia coli*^28,29^ and *Komagataella phaffii*^30^. Thus, proving the feasibility of synthetic CO_2_ fixation pathways in vivo remains an open challenge. In vivo validation is essential to determine real-life performance in the context of the cellular metabolic network and to compare pathways and modules to discover the most promising ones.

Here, we propose new pathway architectures based on crotonyl-CoA carboxylase/reductase (CCR) and 2-oxoglutarate carboxylase (OGC), which we name reductive methylaspartate (rMASP) cycles. We show that this architecture lends itself to several variants with different properties and products and use a modular engineering approach to divide each pathway into modules, which we test in vivo in *E. coli* with growth coupled-selection strains where the end product of each module relieves a specific auxotrophy. The rMASP cycles are predicted to be more energy efficient than other synthetic CO_2_ fixation pathways at a comparable pathway activity. Moreover, their overlap with existing designs means that engineering efforts towards the rMASP cycle will also be useful towards establishing other synthetic CO_2_ fixation pathways such as the CETCH^18,19^, rGPS-MCG and THETA^21^ cycles.

## Results

### Design and analysis of the reductive methylaspartate cycles

For the design of the rMASP cycle we considered the carboxylation of the C5 molecule 2-oxoglutarate to isocitrate, a reaction that has been underexplored so far in synthetic CO_2_ fixation^14,17^. This can be achieved either via the reverse activity of isocitrate dehydrogenase, which is thermodynamically challenging and requires an elevated CO_2_ atmosphere^31,32^, or via a fast biotin-dependent enzyme, which uses the energy from ATP hydrolysis to drive the reaction and can operate at ambient CO_2_^33–35^. In both cases, the carboxylation product is oxalosuccinate, which is then reduced to isocitrate by an oxalosuccinate reductase using NADH^36^. We then used a retrosynthetic approach to design full oxygen-tolerant pathways around this reaction, resulting in two different pathway architectures that convert the C6 molecule isocitrate into a C2 product (acetyl-CoA or glyoxylate) and the C4 pathway intermediate crotonyl-CoA. (Fig. 1). Crotonyl-CoA carboxylase (CCR), one of the fastest carboxylases known in nature, catalyzes the second CO_2_ fixation step in all pathway variants. The product of this carboxylation, ethylmalonyl-CoA is converted to glutamate via a new-to-nature pathway module, which was not proposed before for synthetic CO_2_ fixation. Glutamate can be deaminated to regenerate 2-oxoglutarate and start another cycle. We analyzed two variants for each of these two architectures, calculating their max-min driving force (MDF)^37^, their theoretical biomass yield using flux balance analysis (FBA)^38^, and their expected pathway activity based on the enzyme cost minimization (ECM)^39^ algorithm (Fig. 2 and supplementary Table 1). We compared the rMASP cycles to the CBB cycle considering both a high CO_2_ scenario and an ambient CO_2_ scenario. We also included a comparison to other proposed synthetic CO2 fixation pathways (CETCH^18^, GED^40^, HOPAC^41^, rGPS-MCG^20^, THETA^21^). We divided each of the four resulting cycles into three modules, and used a modular engineering approach to implement them in vivo. We named each module after its output molecule: glutamate, isocitrate, and crotonyl-CoA. The glutamate and isocitrate modules are constant, whereas the crotonyl-CoA module has four variants, named after their characteristic enzymes isocitrate lyase (ICL), ATP-citrate lyase (ACL), acetoacetyl-CoA thiolase (AACT), and acetoacetyl-CoA synthase (AACS) (Figure 2A). In this paper, we report the full in vivo implementation of the glutamate module, the partial in vivo implementation of the crotonyl-CoA modules, and the in vitro demonstration of the isocitrate module.

**Figure 1.**
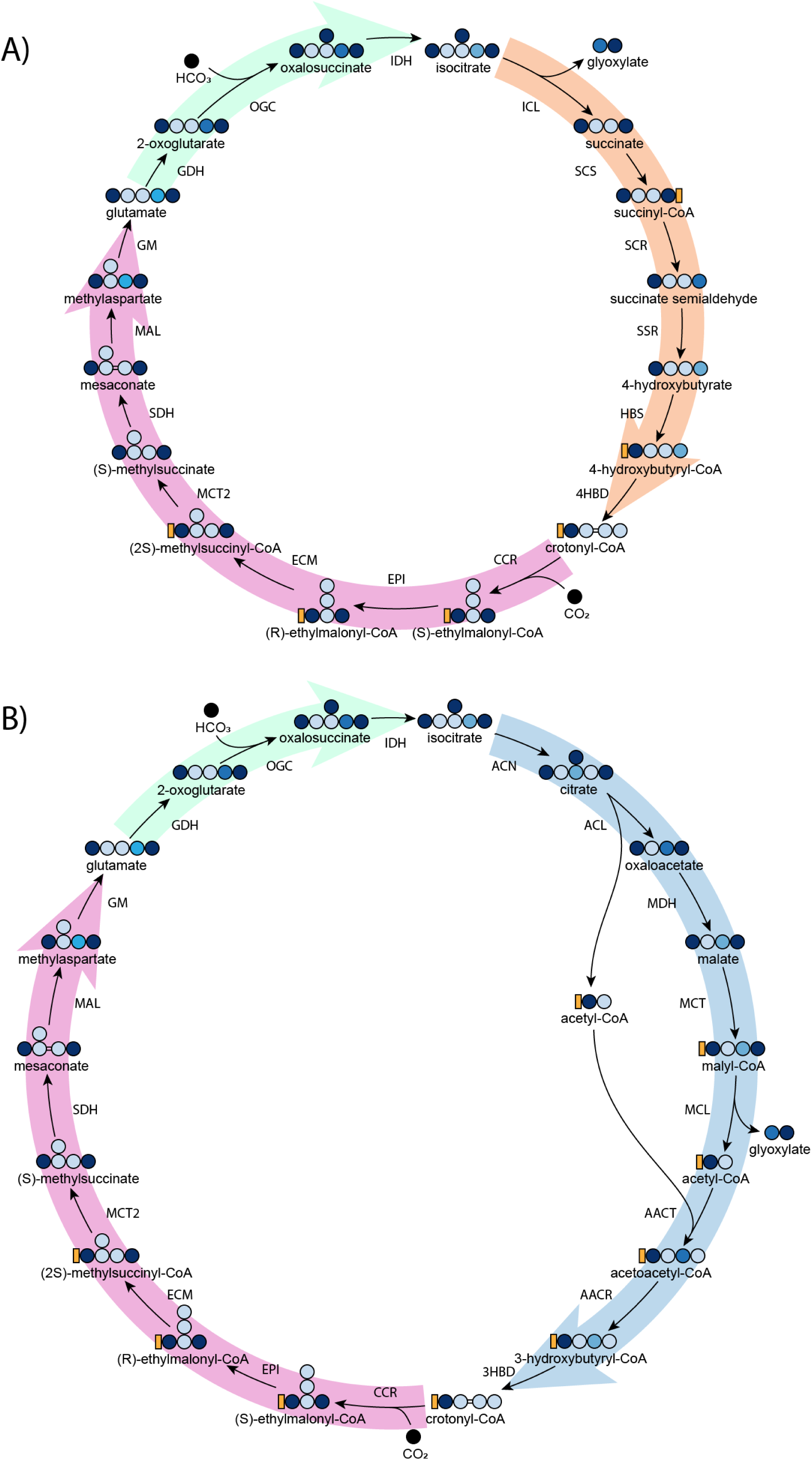
The two different pathway architectures of the reductive methylaspartate cycle. (A) via isocitrate lyase (ICL) and (B) via acetoacetyl-CoA thiolase (AACT). 3HBD, 3-hydroxybutyryl-CoA dehydratase; 4HBD, 4-hydroxybutyryl-CoA dehydratase; ACL, ATP citrate lyase; ACN, aconitase; AACR, acetoacetyl-CoA reductase; AACT, acetoacetyl-CoA thiolase; CCR, crotonyl-CoA carboxylase; ECM, ethylmalonyl-CoA mutase; EPI, ethylmalonyl-CoA epimerase; ICL, isocitrate lyase; IDH, isocitrate dehydrogenase; GDH, glutamate dehydrogenase; GM, glutamate mutase; HBS, 4-hydroxybutyryl-CoA synthase; HBD, 4-hydroxybutyryl-CoA dehydratase. MAL, mesaconate ammonia lyase; MCL, malyl-CoA lyase; MCT, malate CoA transferase; MCT2, mesaconyl-CoA transferase; MDH, malate dehydrogenase; SDH, methylsuccinate dehydrogenase; OGC, 2-oxoglutarate carboxylase; SCS, succinyl-CoA synthetase; SCR, succinyl-CoA reductase; SSR, succinate semialdehyde reductase.

**Figure 2.**
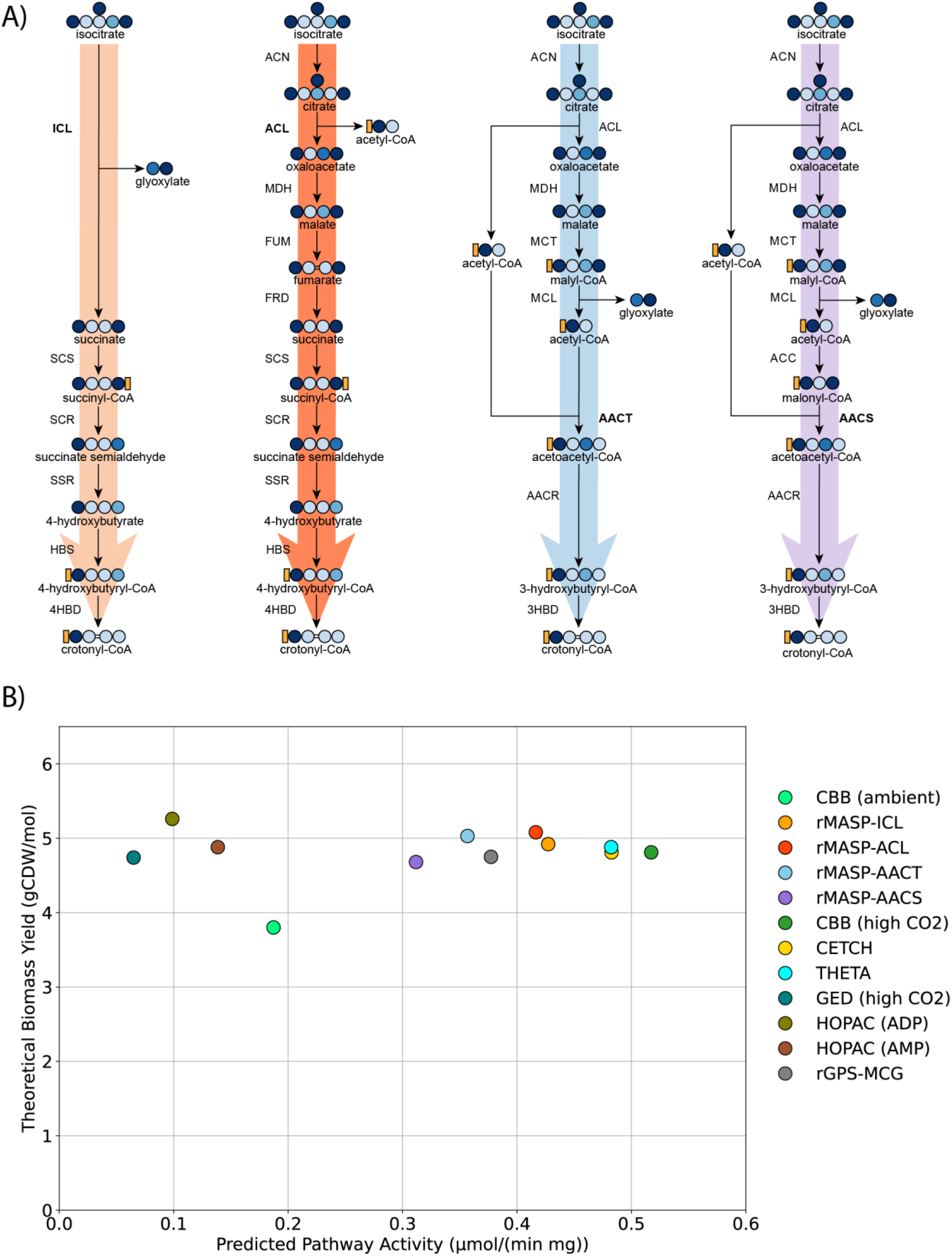
(A) Four modules to regenerate crotonyl-CoA from isocitrate (B) Computational prediction of biomass yield (calculated using FBA) and pathway activity (calculated using ECM) for each variant. Yields are calculated per mol NADH. The ambient CO_2_ condition for the CBB assumes a 20% rate of rubisco oxygenation^42^. AACS, acetoacetyl-CoA synthase; ACC, acetyl-CoA carboxylase; FRD, fumarate reductase; other enzyme abbreviations are the same as in figure 1.

#### In vivo implementation of the glutamate module

The glutamate module is shared across all cycle variants and was never demonstrated in vitro or in vivo (Figure 1, pink arrows). For this reason, we used it as the starting point for in vivo implementation in a glutamate auxotroph strain.

In this work, we used an *E. coli* isocitrate dehydrogenase (Δ*icd*) knockout strain from the KEIO collection^43^. This strain is unable to synthesize 2-oxoglutarate (2OG) and hence is auxotrophic for glutamate and glutamate-derived amino acids. This strain can only grow on glycerol minimal medium with 2OG or glutamate supplementation (figure 3A). Using this strain, we tested the module in a stepwise manner, starting from the conversion of methylaspartate to glutamate, a reaction catalyzed by the B_12_-dependent enzyme glutamate mutase.

**Figure 3.**
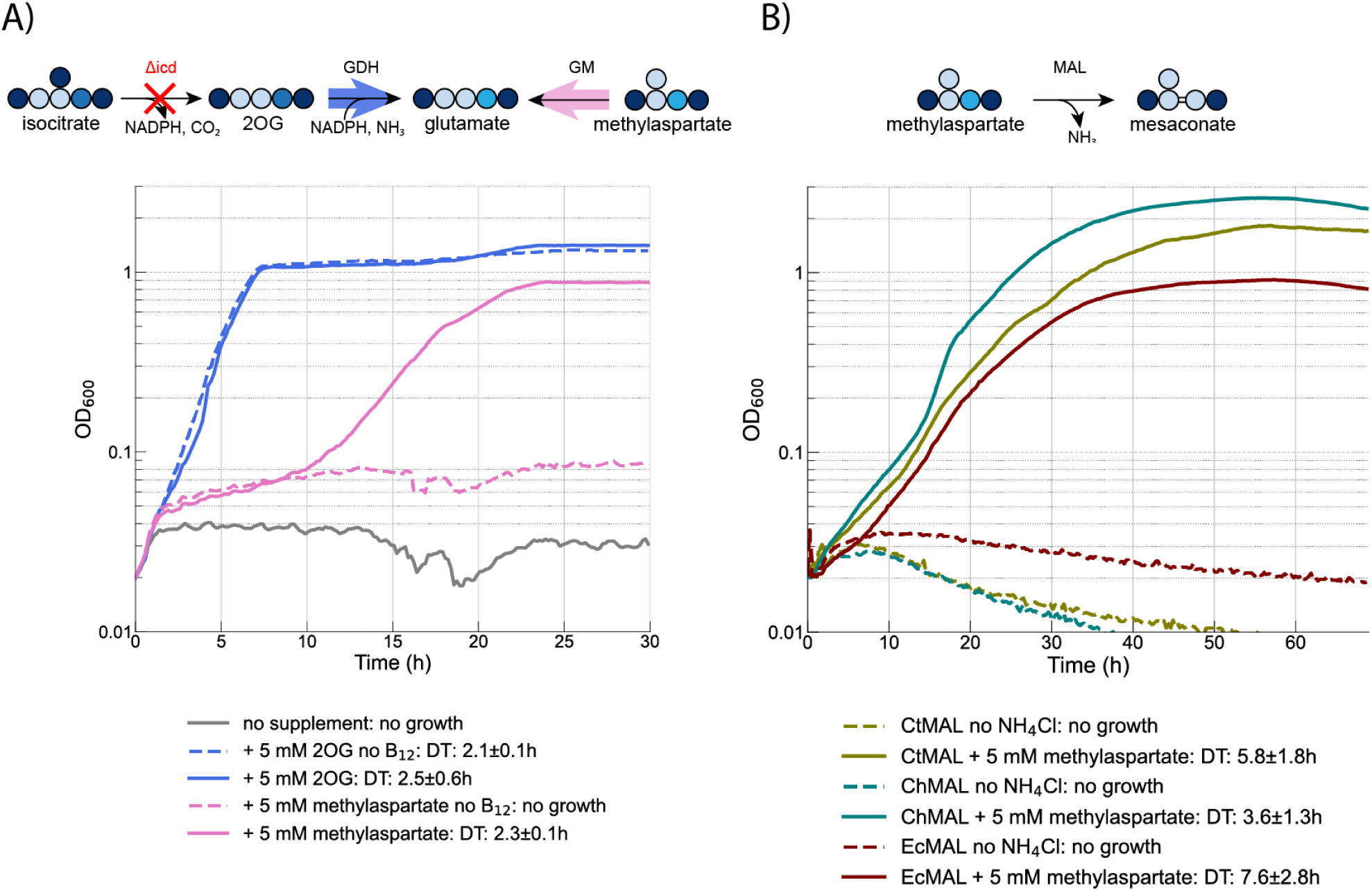
(A) Growth of an *E. coli* glutamate auxotroph (Δ*icd*) via B_12_-dependent glutamate mutase *sanUV* from *S. ansochromogenes*. The strain requires both methylaspartate and B_12_ to produce glutamate. The strain was grown in M9 minimal medium with 20 mM glycerol as a main carbon source and supplements as indicated in the legend. All experiments were performed under fully aerobic conditions, demonstrating that the chosen glutamate mutase is not inactivated by oxygen. (B) In vivo demonstration of functional mesaconate ammonia lyase (MAL) expression. The strains were grown in M9 minimal medium devoid of a nitrogen source (M9N0). Supplementation with methylaspartate allows growth via deamination to mesaconate. Growth curves are the average of at least three technical replicates. DT, doubling time; GDH, glutamate dehydrogenase; GM, glutamate mutase; MAL, methylaspartate ammonia lyase.

For this reaction, we tested several enzyme variants from different organisms that were previously reported to be functional in vitro under anaerobic conditions when purified from *E. coli*^44,45^, but ultimately none of these allowed aerobic growth. We then selected an enzyme from *Streptomyces ansochromogenes*^46^. Since *Streptomyces* spp. are obligate aerobes, the enzyme isoform was expected to be oxygen-tolerant. When we expressed the two genes *sanU* and *sanV* from a plasmid, we observed growth in the Δ*icd* strain using glycerol as a main carbon source and methylaspartate as precursor to generate glutamate. The activity of glutamate mutase is further confirmed by the absence of growth when the coenzyme B_12_ precursor cyanocobalamin is not added to the growth medium, since *E. coli* cannot synthesize it^47^ (figure 3A).

The growth phenotype also indicates that the enzyme is sufficiently fast and that methylaspartate is imported into the cell, presumably via one of the C4-dicarboxylate transport systems such as DctA or DcuA^48^. To the best of our knowledge this is the first reported evidence of glutamate mutase activity in the direction of glutamate formation for this thermodynamically reversible reaction (ΔrG’^m^ = -7.0 ± 4.6 kJ/mol).

After showing that glutamate mutase is functional in vivo, we tested the next step in the module, which produces methylaspartate from mesaconate and ammonium via the activity of methylaspartate ammonia lyase (MAL). This reversible reaction is akin to aspartate ammonia lyase, encoded in *E. coli* by the gene *aspA*, hence we first tested whether this native enzyme could promiscuously accept mesaconate. Since we already showed that methylaspartate is imported into the cell, and since its deamination produces ammonium, we first tested the enzymatic activity in the deamination direction in a wild-type strain using a medium devoid of a nitrogen source. The absence of growth in this experiment suggests that native aspartate ammonia lyase has no or insufficient activity with methylaspartate and hence cannot support growth (figure S1). At this point, we chose three MAL enzyme variants for heterologous expression, originating from *Carboxydothermus hydrogenoformans*^49^, *Clostridium tetanomorphum*^50^, and the pathogenic *E. coli* strain O157:H7^51^. All three enzymes were able to produce ammonium from methylaspartate in vivo, supporting growth in minimal medium without nitrogen addition (figure 2B).

This result still leaves the question of activity in the amination direction required for the CO_2_ fixation pathways, which was previously shown in vitro^49^, but not yet in vivo. To test this activity, the Δ*icd* strain was used once again, this time expressing both glutamate mutase and MAL. We selected ChMAL since this gene performed best in the deamination direction (Figure 3B). We expected this strain to grow on glycerol with mesaconate supplementation, however we did not observe growth. Since we showed methylaspartate ammonia lyase activity in the deamination direction, incorrect expression or folding can be ruled out, however the enzyme could be affected by product inhibition or allosteric regulation by other metabolites. Additionally, mesaconate could be diverted to a different pathway, or it may not be imported well into the cells, thereby limiting the intracellular concentration and decreasing the MAL reaction rate. To solve these issues, we started an adaptive laboratory evolution (ALE) experiment, feeding the strain with glycerol, a limiting amount of 2-oxoglutarate, and an excess of mesaconate (10 mM) and ammonium chloride (100 mM) to increase the thermodynamic driving force of the reaction. Under these conditions, there is a selective pressure for mesaconate consumption, so cells that accumulate beneficial mutations are able to grow more and eventually overtake the population. After about three weeks of passaging, the strains could grow reliably without the addition of 2-oxoglutarate.

At this point we decided to test whether methylsuccinate could also relieve the auxotrophy in the mesaconate evolution strain, since *E. coli* succinate dehydrogenase (Sdh) has previously been shown to convert S-methylsuccinate into mesaconate in vitro^21^. Initially, growth was once again slow, but after subjecting the cells to short term evolution by two weeks of passaging a strain emerged that could use methylsuccinate to relieve the glutamate auxotrophy (figure 4). This strain, which we named V1, was isolated and sent for whole genome sequencing. The observed mutations are reported in Supplementary Table 2. From the observed mutations, *sdhA* F119L seems highly relevant for the observed growth phenotype, since methylsuccinate is not the preferred substrate of wild-type Sdh, and succinate is likely always present intracellularly at mM concentrations, creating a competition between the two substrates. The F119L mutation is in the active site of the enzyme, where F119 seemingly coordinates one of the aliphatic carbons of succinate (Figure S2). The substitution of the hydrophobic amino acid phenylalanine with a smaller, also hydrophobic amino acid could create additional space for the extra methyl group in methylsuccinate, thereby increasing affinity.

**Figure 4.**
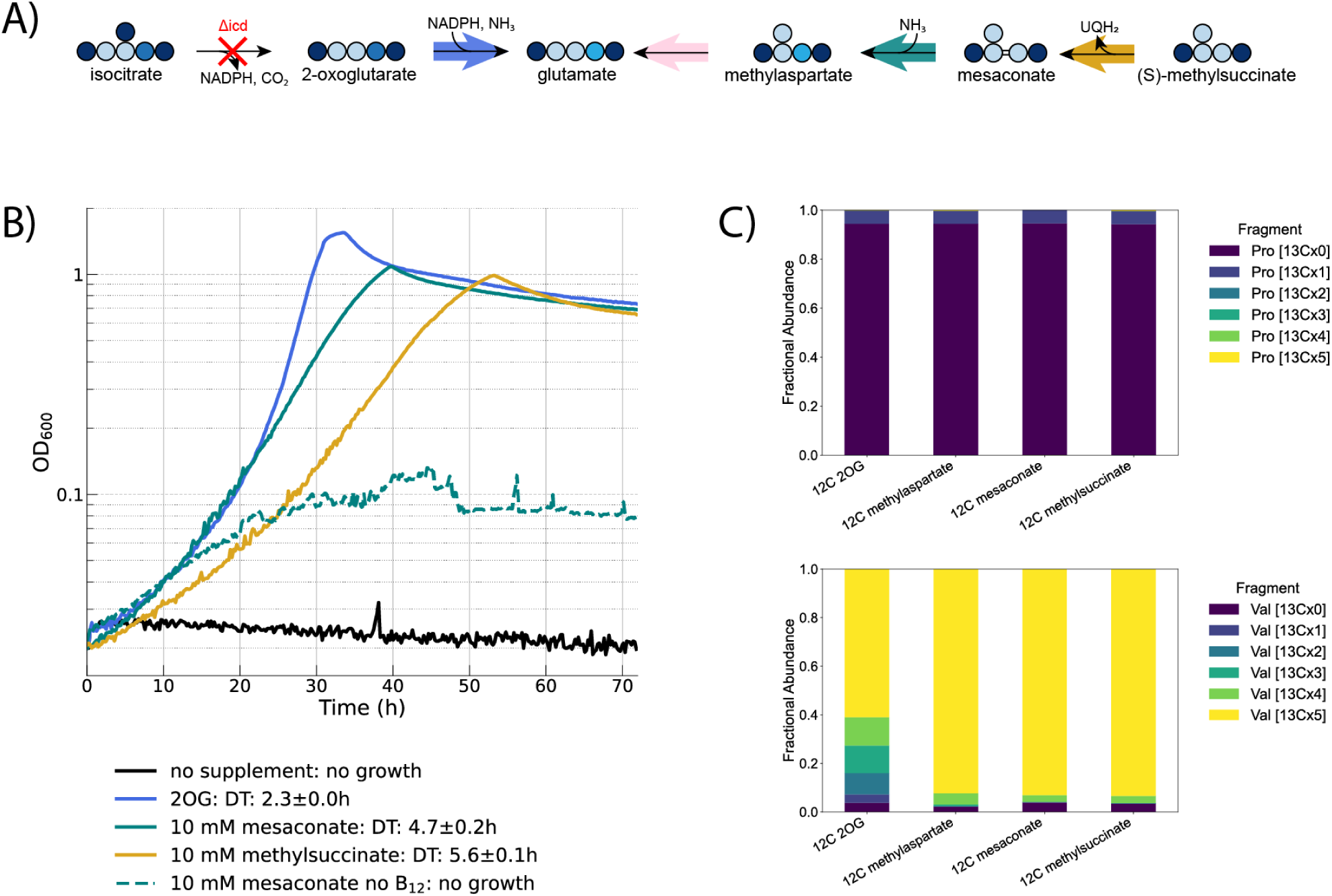
(A) Metabolic scheme of the growth of an *E. coli* glutamate auxotroph (Δ*icd*) via a B_12_-dependent glutamate mutase. The strain cannot grow using glycerol as a sole carbon source, and requires supplementation of a source of glutamate. (B) Growth curves when different supplements are added. Growth is B_12_-dependent, as indicated by the no growth phenotype when B_12_ is absent from the medium (dashed grey line) (C) Isotopic labeling data showing the incorporation of unlabeled carbon sources into biomass. All strains were grown in M9 with 20 mM fully ^13^C labeled glycerol and unlabeled (^12^C) supplementary carbon sources as indicated in the figure. Proline is derived from glutamate and hence expected to be fully unlabeled, whereas valine is derived from pyruvate and expected to be fully labeled. The 2OG condition shows that this metabolite is partially converted to pyruvate via the TCA cycle.

To demonstrate pathway activity, we performed an isotopic labeling experiment to trace the provenance of glutamate in various growth conditions. Since methylaspartate, mesaconate, and methylsuccinate are relatively uncommon molecules, they are not easily available as ^13^C-labeled chemicals. For this reason, we decided to perform a “reverse” labeling experiment, where a main carbon source, in this case ^13^C-glycerol, is added to the medium, while the supplementary carbon source, from which the metabolite of interest is derived, is unlabeled. In this condition, we expected glutamate and glutamate-derived amino acids to be fully unlabeled, and other amino acids (derived from glycerol) to be fully labeled. Our experimental data matches the expectation, as shown in the labeling pattern of proline and valine (Figure 4C). The full labeling data is available in supplementary data file 1.

After achieving methylsuccinate-dependent growth, we tested the remainder of the mutase module for the carboxylation of crotonate and conversion to methylsuccinate. This was previously tested in vivo as part of the THETA cycle, but it could not support growth^21^. We decided to use our evolved glutamate auxotroph strain to test the same constructs. For this reason, we first modified the strain for genomic expression of heterologous ethylmalonyl-CoA mutase (ECM) and epimerase (EPI) and then expressed a crotonate thiokinase (CTK) and a crotonyl-CoA carboxylase (CCR) from an IPTG-inducible plasmid. With the addition of these four enzymes, we expected the strain to be able to use crotonate to relieve the glutamate auxotrophy. Unfortunately, the strain did not grow regardless of IPTG inducer concentration or carbon source tested. For this reason, we decided once again to start an ALE experiment, this time using gluconate as a main carbon source, a limiting amount of methylsuccinate, and an excess of crotonate. We chose gluconate because its degradation via gluconate dehydrogenase (*gnd*) can produce stoichiometric amounts of NADPH, which is the source of reducing power for CCR. Additionally, we added bicarbonate to increase the amount of CO_2_ available for the carboxylation reaction. After about one month of passaging, a strain emerged that could grow without methylsuccinate. We isolated and sequenced this strain, which we named V2 and presents several mutations compared to its parent (Figure S3, Supplementary Table 2). Even though CCR is reported to operate at ambient CO_2_ concentrations, we only observed minimal growth without extra CO_2_ in the headspace or the addition of bicarbonate to the growth medium. So, we could demonstrate full growth-supporting activity of the glutamate module, including crotonyl-CoA carboxylation, but not yet at ambient CO_2._

#### Partial in vivo implementation of crotonyl-CoA regeneration modules

After the implementation of the glutamate module, we tested the crotonyl-CoA regeneration modules by curing the plasmid containing CTK and CCR (pTE3288), obtaining the V3 strain, which we used to test a different plasmid encoding all genes necessary for succinate conversion to crotonyl-CoA (pTE3236C). In this plasmid, CTK is replaced by a crotonyl-CoA transferase (CCT), which was previously shown to have superior kinetic parameters on both crotonate and 4-hydroxybutyrate in vitro^21^. Since all required enzymes are present, we expected this strain to grow by using succinate to produce glutamate, however we did not observe growth (Fig. 5A). We confirmed that growth on crotonate was still possible, however we observed that addition of succinate in the growth medium abolished crotonate-dependent growth. This could be attributed to competitive inhibition for Sdh activity between succinate and methylsuccinate. Since the latter activity is essential for growth on crotonate, its inhibition could slow down or even abolish growth. We also tested supplementation with 4-hydroxybutyrate, which requires the additional activity of 4-hydroxybutyryl-CoA dehydratase compared to crotonate, however we did not observe growth. Since in vivo activity was previously confirmed for all enzymes converting succinate to crotonyl-CoA^21^, a lack of growth could be attributed to backwards flux from 4-hydroxybutyrate to succinate, since the pTE3236C plasmid encodes for NAD-dependent dehydrogenases for succinyl-CoA and succinate semialdehyde^21^, which are likely thermodynamically constrained in the reductive direction. To test this hypothesis, we recloned the plasmid to only include the dehydratase, CoA transferase, and carboxylase encoding genes (HBD-CCT-CCR, pTE3236D). This time, we observed 4-hydroxybutyrate-dependent growth when the plasmid was induced, indicating its conversion to glutamate (Fig. 5B).

**Figure 5.**
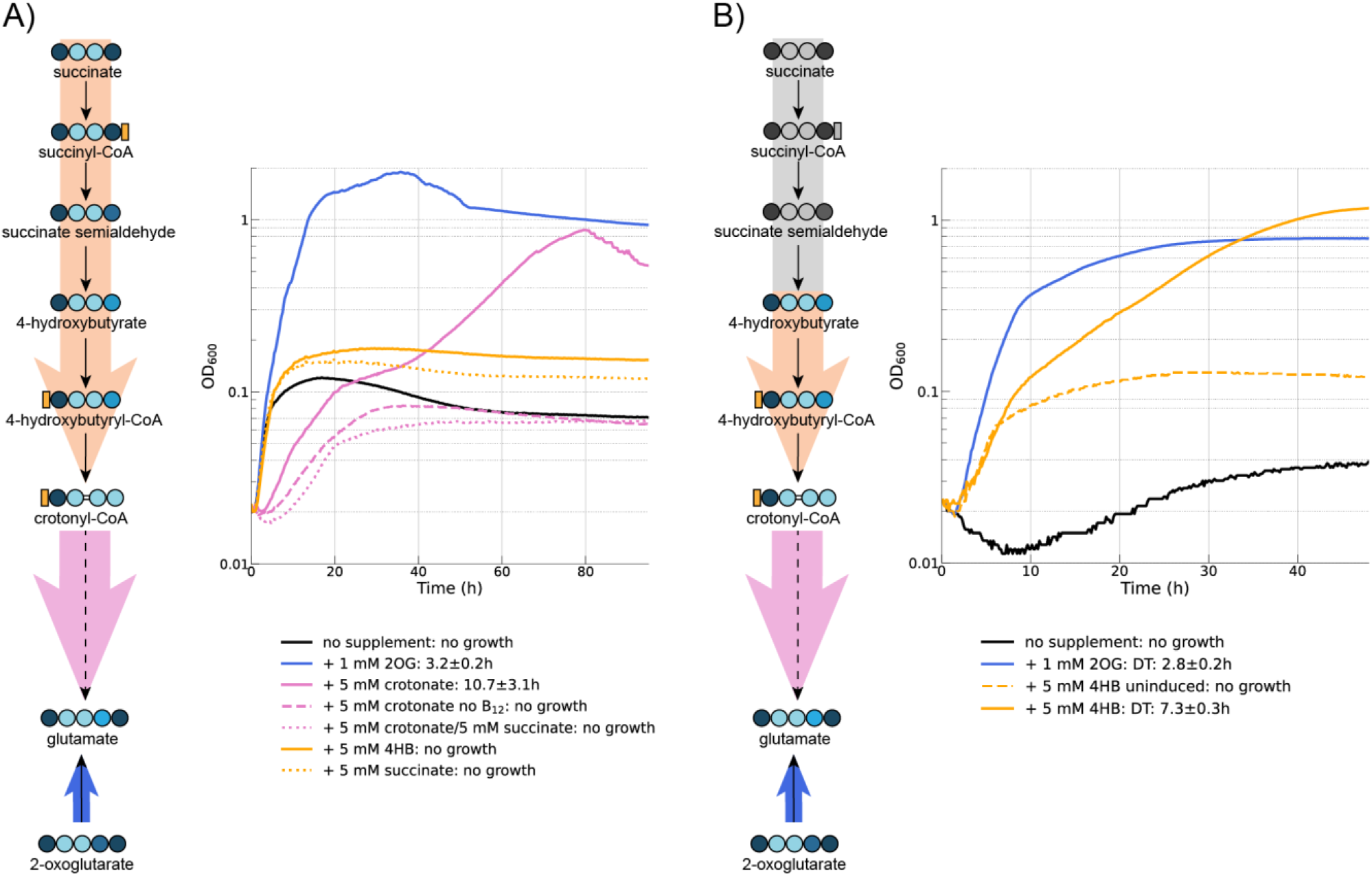
**(A)** Crotonate-dependent growth of the V3 strain carrying pTE3236C via the full mutase module (solid pink line). Addition of succinate (dotted pink line) inhibits growth, potentially indicating a toxicity effect due to the competition between succinate and methylsuccinate for Sdh activity. Neither succinate nor 4HB (orange lines) support growth in the absence of crotonate **(B)** 4HB-dependent growth of the V3 strain carrying pTE3236D, in which the greyed out enzymes were removed. Experiments were performed in M9N100 with 10 mM gluconate, 10 µM B_12_, 50 µM IPTG, 5% CO_2_ and supplemental carbon sources as indicated in the legend. B_12_ or IPTG were omitted as negative controls, indicated by dashed lines. 2OG, 2-oxoglutarate; 4HB, 4-hydroxybutyrate; DT, doubling time

Next, we tried to implement a different crotonyl-CoA regeneration module variant based on the first part of the ethylmalonyl-CoA pathway (EMCP) to regenerate crotonyl-CoA from two acetyl-CoA molecules, bypassing the need for succinate (Fig. 6). There are two variants of this module, one of which uses the native *E. coli* acetoacetyl-CoA thiolase (AACT, encoded by *atoB*), while the other one uses the heterologous acetoacetyl-CoA synthase (AACS, encoded by *nphT7* in *Streptomyces sp. CL190*).

**Figure 6.**
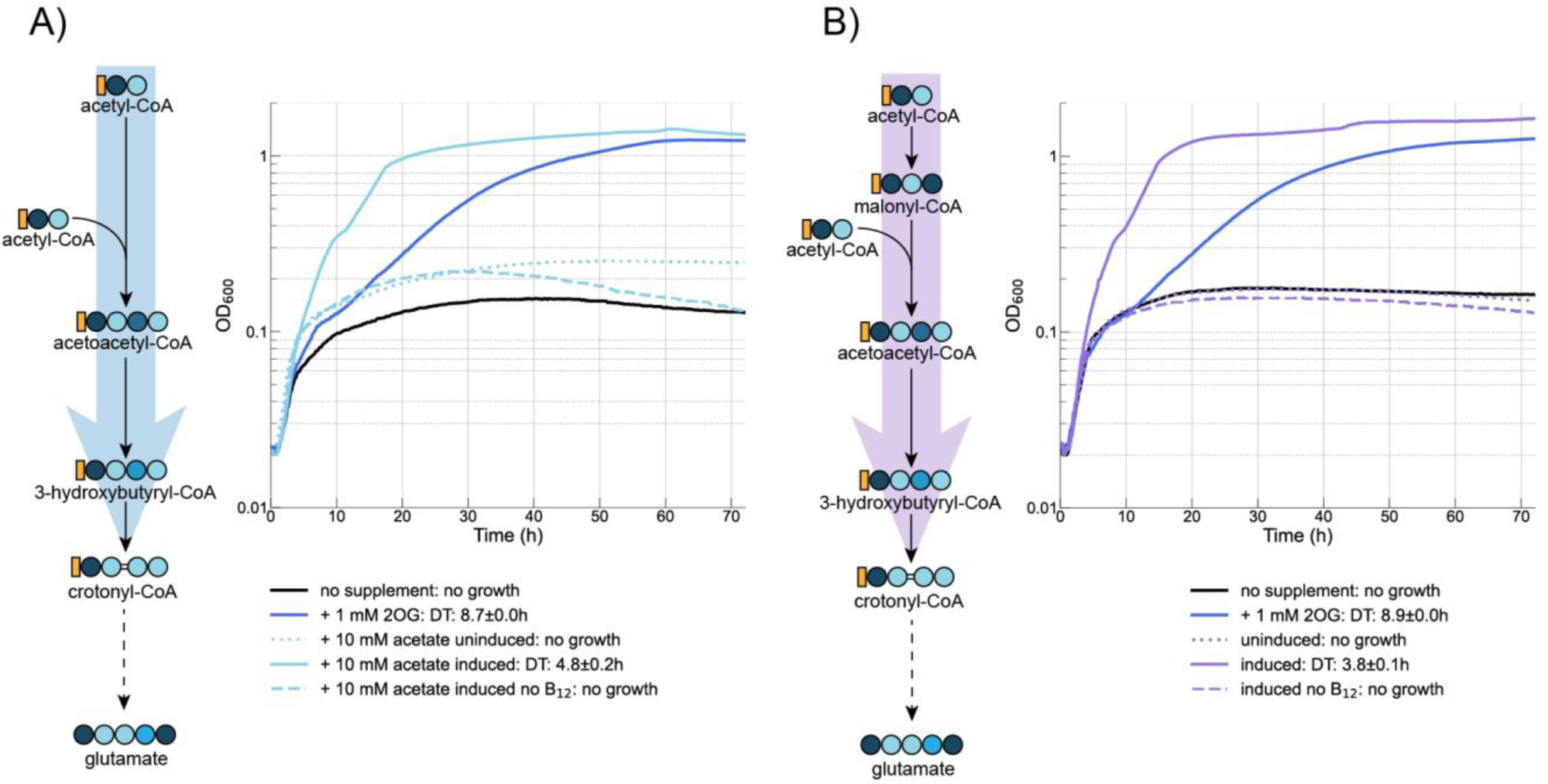
Growth of the V3 strain via the AACT (left) and AACS (right) modules for acetate assimilation. Strains were grown in M9N100 with 10 mM gluconate as a main carbon source, 10 µM B_12_, 10 µM cuminic acid, 5% CO_2_ and supplements as indicated in the legend. Dashed or dotted lines indicate the omission of B_12_ or cuminic acid, respectively.

The former is more energetically efficient, while the latter benefits from additional thermodynamic driving force^52^. Expression of either variant from a plasmid with a cumate-inducible promoter led to growth (Fig. 6). Notably, the AACS variant did not require acetate supplementation in the growth medium, presumably because the acetyl-CoA produced during gluconate degradation is sufficient for pathway activity, whereas the AACT variant may require a higher acetyl-CoA concentration to achieve sufficient thermodynamic driving force for the reaction. Furthermore, both modules currently use the native *E. coli* 3-hydroxyacyl-CoA dehydrogenase FadB which is NAD-dependent and hence thermodynamically less favorable in the reductive direction. This enzyme is normally involved in fatty acid degradation (hence the name), and operates in the oxidative direction, producing NADH. Swapping this enzyme for an NADPH-dependent isozyme, e.g. from *Cupriavidus necator*^52^ will improve driving force and kinetic parameters for both variants.

The demonstrated parts of rMASP cycles via the AACT and AACS variants constitute in total enzyme cascades of 10 and 11 enzymes respectively, demonstrating a major part of the total rMASP cycles (18-19 enzymes). On their own, both routes can also serve for growth starting from acetate and bioproduction of e.g. the valuable chemical glutamate. This is especially relevant as acetate is a promising sustainable feedstock for future biotechnology^53^. In nature, there are only three known pathways for growth on acetate: the glyoxylate cycle^54^, the EMCP^55^, and the methylaspartate cycle of Haloarchea^56^. Additionally, a reductive citramalyl-CoA shunt was proposed but never demonstrated^16^. Our computational flux balance analysis shows that the AACT variant is predicted to be the most efficient for growth on acetate, and for glutamate production from acetate, outperforming all known natural routes (Supplementary Table 3).

### The CCR from *Methylorubrum extorquens* allows growth at ambient CO_2_ concentration

Throughout this work, we used the CCR from *R. sphaeroides* (RsCCR) in all our constructs, since this variant was previously used in vitro test and in vivo attempts of the implementation of the THETA cycle. This enzyme is the best characterized CCR, but it shows relatively poor kinetic parameters with regards to affinity for both NADPH and CO_2_ (0.7 mM and 0.2 mM, respectively). The concentration of NADPH in *E. coli* is known to be around 0.1 mM^57^ and the concentration of dissolved CO_2_ at atmospheric pressure is between 10 and 20 µM^58^, meaning that the enzyme is likely operating well below its maximal velocity under ambient CO_2_ conditions. We reasoned that this likely resulted in the observed lack of growth observed without high CO_2_ partial pressure or bicarbonate supplementation. Since functionality at atmospheric CO_2_ concentrations is a core criterion for synthetic CO_2_ fixation in photosynthetic conditions with ambient CO_2_ and O_2_, we decided to test an alternative CCR from *Methylorubrum extorquens* (MeCCR). While the affinity for CO_2_ of this specific isozyme has not been reported, we expected better performance, since *M. extorquens* grows via the EMCP using the serine cycle in an environment (the phyllosphere) where CO_2_ concentration is not higher than normal, and in fact could even be CO_2_ depleted due to the photosynthetic activity of leaves. By substituting RsCCR with MeCCR we observed 4HB-dependent growth in ambient cultivation conditions (Fig. 7). Notably, the doubling time improved from 7.3 h to 5.7 h, indicating that MeCCR is kinetically faster at ambient CO_2_ than RsCCT at high CO_2_. Furthermore, there is no difference in doubling time between MeCCR at ambient CO_2_ vs high CO_2_ (5.7 h vs 5.4 h, fig. S5) which indicates that ambient CO_2_ is likely saturating.

**Figure 7.**
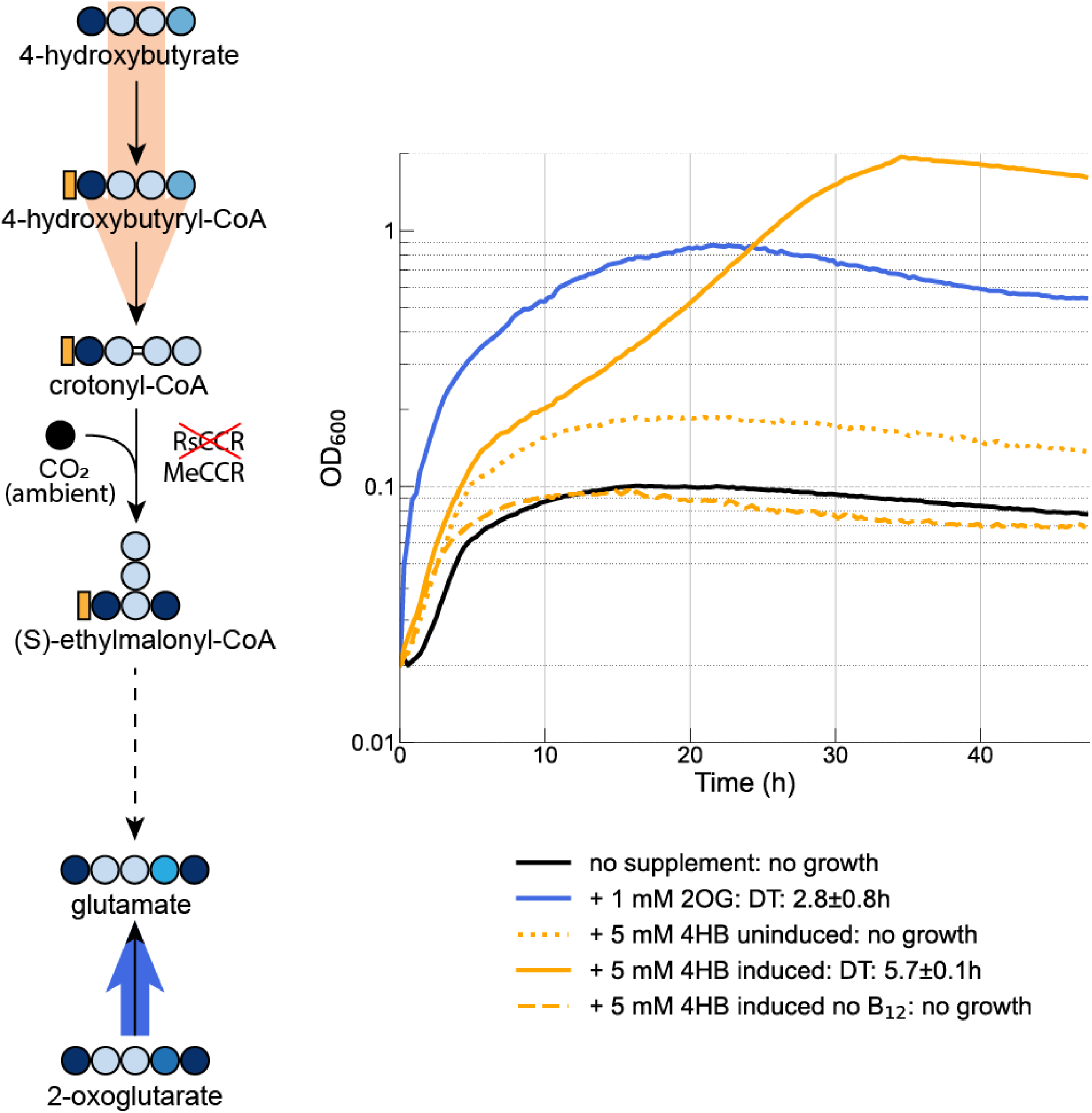
Growth of the V3 strain carrying pTE3236E in M9N100 with 10 mM gluconate, 10 µM B_12_, and 50 µM IPTG, and supplementary carbon sources as indicated in the legend under ambient CO_2_ concentration. Dotted and dashed lines indicate the omission of IPTG or B_12_ from the medium, respectively. 2OG, 2-oxoglutarate; 4HB, 4-hydroxybutyrate; DT, doubling time.

#### In vitro testing of the 2OG carboxylation module

The last module of the rMASP cycles produces isocitrate from the carboxylation of 2-oxoglutarate via OGC from *Hydrogenobacter thermophilus*. While this enzyme has been heterologously expressed in *E. coli* and assayed in vitro, this assay was based on ATP hydrolysis, and hence not coupled to product formation^35^. In *H. thermophilus*, the product of the OGC reaction, oxalosuccinate, is reduced to isocitrate via an NADH-dependent, non-decarboxylating ICD. The same reactions are part of the isocitrate module in the rMASP cycles. We heterologously expressed HtOGC and HtIDH in *E. coli* and performed a coupled assay where NADH consumption can be monitored at 340 nm, confirming the activity of both enzymes at 60 °C (Fig. 8, supplementary data file 2). Previous work reported that the temperature optimum for HtOGC is 78 °C, and that lower temperatures negatively affect kinetics, with about half the activity at 48 °C^29^. We tested the intermediate temperature of 37 °C, since this is the optimum growth temperature of *E. coli*, and found no activity. Notably, even at 60 °C our results are much worse than reported values (K_M_ 1.03 mM, V_max_ 14.6 at 70 °C)^33^. The same authors also report activity at 20 °C^33^. The cause of this discrepancy is unknown, however it is possible that expression or folding issues in our system hamper the activity of the enzyme. Lack of activity at 37 °C could be attributed to OGC, IDH, or both. For this reason we tested IDH alone and found that it could catalyze the oxidation of isocitrate to oxalosuccinate at 37 °C, albeit at a relatively poor rate (supplementary data file 3). This result demonstrates that OGC is likely inactive at 37 °C, which is consistent with the fact that this enzyme is complex and undergoes significant conformational changes during catalysis. This shows that for successful in vivo implementation of this third module under mesophilic temperature conditions, enzyme engineering and/or identification of so for unknown mesophilic variants of the IDH and OGC enzymes is needed.

**Figure 8.**
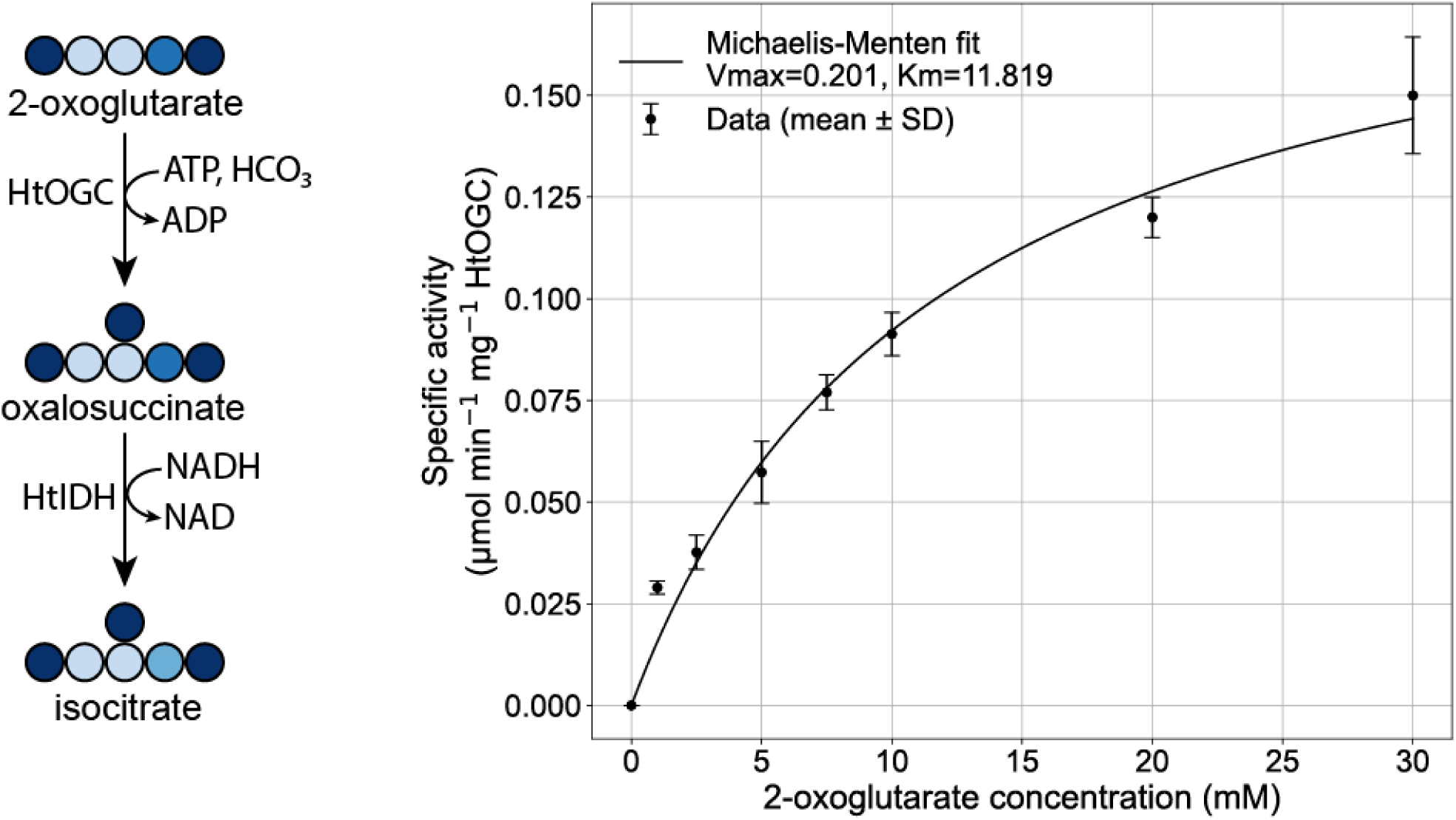
In vitro coupled assay of HtOGC and HtIDH after heterologous expression in *E. coli* and purification. The assay was performed at 60 °C in triplicate.

## Discussion

In this study, we presented the novel rMASP CO_2_ fixation pathways, which have a higher theoretical biomass yield than other natural and synthetic counterparts, such as the CBB, CETCH, THETA and rGPS-MCG cycles. Our design represents a new, previously unexplored family of carbon fixation cycles based on an uncommon carboxylase activity found in hyperthermophilic chemolithotrophs. Until recently, none of the proposed synthetic pathways used this enzyme, showing that new carboxylation reactions can unlock entirely new modes of carbon fixation with the potential to be more efficient than current alternatives. Notably, a comprehensive computational analysis of all possible carbon fixation pathway based on known biochemical reactions recently failed to identify the rMASP cycle^17^. This can be attributed to the fact that the promiscuous activity of succinate dehydrogenase on methylsuccinate is not annotated in databases and hence was not included in that work. This highlights the need for curation of reaction sets based on biochemical expertise when performing such computational analyses.

Here, we divided our proposed pathway into three metabolic modules and showed that two of them are functional in vivo based on growth-coupled selection strains. While some pathway modules have previously been tested and shown to work in vitro, our modular engineering approach, coupled with short-term evolution, was instrumental in finding solutions to unexpected challenges that only arise in vivo, such as the presence of alternative sinks.

More work will be necessary to fully establish the pathway by testing all enzymes in each module and eventually combining them in the same strain while reducing or eliminating current bottlenecks such as slow enzyme kinetics. Additionally, the glutamate module is currently operating in a way that is energetically suboptimal, relying on a promiscuous thioesterase to cleave the CoA from methylsuccinyl-CoA to obtain methylsuccinate, leading to a loss of ATP^21^. This could be prevented in the future by finding or engineering a CoA ligase or transferase that is specific for this substrate^59^. Alternatively, the promiscuous activity of succinate dehydrogenase on methylsuccinate could be substituted by a heterologous methylsuccinyl-CoA dehydrogenase, eliminating potential issues with competitive inhibition due to the presence of succinate in the cell^41,60^. This would require a CoA ligase or transferase that is active on mesaconyl-CoA instead^59^.

Another current challenge of the rMASP cycles is the requirement for a high concentration of ammonium chloride in the medium. This is likely due to poor affinity of the tested MAL from *C. hydrogenoformans* for this co-substrate. While this value has not been reported, the enzyme was used in the amination direction with 400 mM NH_4_Cl^49^. This is likely an issue related to this specific isoform of the enzyme, since similar enzymes such as aspartate ammonia lyase from *E. coli* show much more favorable affinity for NH_4_Cl in the mM range^61^. Additionally, better kinetics for the reactions before and after MAL would provide additional thermodynamic driving force by producing more of the substrate and removing more of the product, respectively.

The lack of activity of OGC at 37 °C will have to be solved to establish the full cycles in vivo. This could be achieved by finding a mesophilic variant of the enzyme, or by engineering a mesophilic pyruvate carboxylase (PYC) to accept the larger substrate 2-oxoglutarate. A database search for mesophilic variants of the enzyme did not yield positive results, however given the similarity between OGC and PYC, the former could be misannotated as the latter. Another option could be engineering HtOGC to lower its temperature optimum and increase activity at mesophilic temperature. Increasing enzyme T_m_ is a common problem in biotechnology, e.g. for the production of thermostable enzymes for detergents or biocatalysis, and much effort has been expended to solve it, while engineering thermophilic enzymes to work at mesophilic temperatures is a relatively unexplored field. Many of the same techniques could be applied, such as error-prone PCR and computer modeling, including machine learning approaches based on datasets of mesophilic vs. thermophilic proteins. More research will be needed to test these approaches and know which one of them will yield results.

Finally, multiple rounds of adaptive laboratory evolution were necessary to achieve growth, first for mesaconate and methylsuccinate, then for crotonate. The strains present numerous mutations compared to the original Δ*icd* strain that we used as the basis for this work. While some of these mutations, such as *sdhA* F119L are relatively easy to interpret and connect to a growth phenotype, others are much less clear. A reverse engineering approach could be applied in the future to transfer mutations stepwise to the original unmutated strain and test which ones are strictly necessary for growth.

The modules implemented in this work have a high degree of overlap with other CO_2_ fixation designs such as the CETCH, rGPS-MCG and THETA cycles. Since these pathways could not yet be engineered in vivo, our evolved strain containing the crotonyl-CoA regeneration and the glutamate modules could serve as starting point for their implementation. The methylaspartate ammonia lyase and glutamate mutase could be replaced with a mesaconyl-CoA hydratase and either citramalyl-CoA lyase or a β-methylmalyl-CoA lyase while reintroducing *icd* as well as adding appropriate deletions rendering the strain auxotrophic for either acetyl-CoA or glyoxylate. While this approach requires extensive modifications of the endogenous metabolism, it would allow exploiting the fact that the difficult to realize reaction sequences of these cycles are already functional in our evolved strain.

Overall, our work represents a major advance over the state of the art of synthetic CO_2_ fixation and the overlap of some metabolic modules with other cycles such as the CETCH and THETA means that lessons from this work will be useful towards the engineering of other pathways, bringing the goal of synthetic CO_2_ fixation closer.

## Materials and Methods

### Materials

All chemicals were purchased from Sigma-Aldrich and Santa Cruz Biotechnology. Fully labeled ^13^C glucose was purchased from Cambridge Isotope Laboratories. Oligonucleotides were purchased from Integrated DNA Technologies (IDT) or Sigma-Aldrich. Synthesized genes were obtained from Twist Bioscience or IDT. All materials for cloning and molecular biology techniques were purchased from New England Biolabs, QIAGEN, and Zymo Research.

### Organic synthesis of 4-hydroxybutyrate

4-hydroxybutyrate was synthesized by base-catalyzed deesterification of gamma-butyrolactone (GBL, purchased from Sigma-Aldrich). 5g GBL was dissolved in 125 ml water, then 2.4g NaOH was added and the reaction was left overnight under reflux at 100 °C. The following day, the reaction mix was cooled to room temperature and the pH was adjusted to 5. After extraction with ethyl acetate (150 mL three times), the organic phase was collected and dried with anhydrous MgSO_4_. A rotary evaporator was used to concentrate the product with a final yield of 90% measured by NMR, using a 500MHz (11.7T) Bruker Avance III NMR spectrometer, with a one-dimensional ^1^H NMR spectrum (zg30, 128 scans) in deuterated DMSO. Data was analyzed with the Bruker TopSpin 4.3.0 software. Attributed spectra are provided in Supplementary Fig. 5.

### MDF and ECM analysis

MDF analysis was applied to evaluate the thermodynamics feasibility of different pathways when considering constrained in vivo metabolite concentrations. Python packages equilibrator_api (version 0.6.0) and equilibrator_pathway (version 0.6.1) were used. The changes in Gibbs energy of the reactions were estimated using the component contribution method. CO_2_ was taken as the substrate for all carboxylation reactions since its concentration is not influenced by pH, unlike that of bicarbonate, thus simplifying the calculations. Ambient CO_2_ in solution is 10-20 μM, high CO_2_ is fixed at 3.4 mM (10 % CO_2_ partial pressure) and other cofactor and metabolite concentrations were constrained to the range 10 μM–10 mM. pH was set to 7.0, ionic strength to 0.25 M and −log[Mg2+] (pMg) to 3.

The pathway activities reported here were determined with enzymatic parameters compiled by Lowe and Kremling^1^, estimated with ENKIE (version 0.1.3)^2^, or sourced from BRENDA. For MDF and ECM calculations we used code adapted from https://gitlab.com/elad.noor/ged-cycle with the equilibrator-cache (0.6.1), equilibrator-api (version 0.6.0) and equilibrator-pathway (version 0.6.0) python package with the following parameters: algorithm ECM, version 3, kcat_source fwd, denominator CM, regularization volume, p_h 7.5, ionic_strength 0.3 molar, p_mg 3, dg_confidence 0.95. The ECM algorithm returns a list of enzyme costs in moles of enzyme per liter based on a total flux through the pathway for 1 mM/s of product formed. These values were multiplied by the molecular weight of the respective enzyme to obtain the enzyme cost in g/l. The sum of all the values is the enzyme cost (EC) for the pathway. To obtain the pathway activity we performed the following calculations:

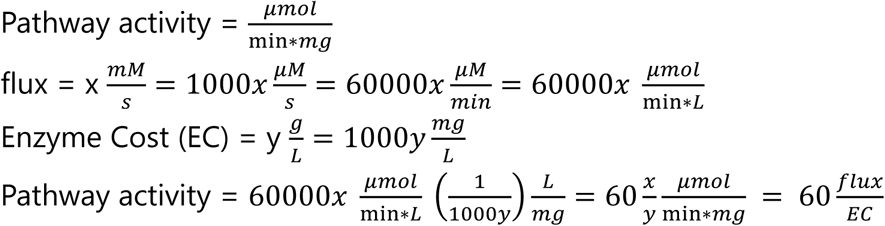

All yield and pathway activity graphs were generated using matplotlib version 3.10.1. Other packages used include numpy (version 2.2.3), pandas (version 2.2.3), tqdm (version 4.67.1).

The scripts can be found on gitlab (https://git.wur.nl/vittorio.rainaldi/r_masp).

### Bacterial strains and growth conditions

*Escherichia coli* DH5α (Thermo Scientific, Dreieich, DE) strains were used for cloning and were grown in LB medium. For protein expression, *E. coli* BL21-AI (Invitrogen, for production of *Ht*IDH, *Ht*OGCS and *Ht*OGCL) auto induction TB medium at an incubation temperature of 25°C. The following antibiotics were used for selection: 35 µg/ml streptomycin and 25 µg/ml kanamycin. *E. coli* strains were routinely grown in LB supplemented with appropriate antibiotics at 37 °C. For growth rescue experiments, M9 minimal medium (8.9 g l^−1^ Na_2_HPO_4_·2H_2_O, 3 g l^−1^ KH_2_PO_4_, 0.59 g l^−1^ NaCl and 1 g l^−1^ NH_4_Cl; pH 7.2), 1 mM MgSO_4_, 0.2 mM CaCl_2_, 50 μM FeSO_4_, trace metals (68.2 μM MnCl_2_, 3.7 μM ZnSO_4_, 0.4 μM CoCl_2_, 0.6 μM CuCl_2_, 1.6 μM H_3_BO_3_, 2.1 μM NiCl_2_) was used with supplementation of appropriate carbon sources as noted in the text. Additional ammonium chloride was added to M9 to reach a final concentration of 100 mM instead of the usual 20 mM. This custom medium is indicated in the text as M9N100. Similarly, M9 devoid of a nitrogen source is referred to as M9N0. Antibiotics were used at the following concentrations: spectinomycin (Spec), 100 μg ml^−1^; kanamycin (Kan), 50 μg ml^−1^; chloramphenicol (Cam), 25 μg ml^−1^; apramycin (Apr), 50 μg ml^−1^; ampicillin (Amp), 100 μg ml^−1^.

### Plasmid construction

All in silico cloning was performed with Geneious (Auckland, NZ) All gene fragments were purchased from GenScript (Rijswijk, NL). All plasmids constructed in this study were assembled using the Golden Gate assembly method. DNA fragments were amplified by Q5 high-fidelity DNA polymerase. Plasmids are listed in Supplementary Table 4. Plasmid maps are available as Supplementary data files. All plasmids were transformed in *E. coli* strain DH5α or DH10β for propagation and storage.

### Strain engineering

Strains used in this study are listed in Supplementary Table 4. λ-Red recombineering was used for gene knockouts and integrations using either the helper plasmid pSIJ8 in the BW25113 strain or its genomically integrated counterpart in the SIJ488 strain^62^. For knockouts, Kan and Cam selection markers were PCR amplified using primers with 50 bp homology arms, from pKD4 (GenBank: AY048743) and pKD3(GenBank: AY048742), respectively. For the genome integration of the epi–ecm operon in SS7^63^, P1 phage transduction^64^ was used from a strain already carrying the integration^21^.

#### Cell growth tests

After transformation or electroporation, single colonies of the strain with the desired plasmids were cultivated overnight in LB with appropriate antibiotics and supplements. On the next day, the overnight culture was washed twice with 0.9% NaCl and inoculated at a starting OD of 0.02 into 3 ml M9 minimal medium with appropriate carbon sources, antibiotics, and inducers in glass tubes. Growth experiments were performed in 96-well microtiter plates (Nunclon Delta Surface, Thermo Scientific) at 37°C and were inoculated with washed cells to an optical density (OD600) of 0.01 in 150 µL total culture volume per well. To avoid evaporation but allow gas exchange, 50 μL mineral oil (Sigma-Aldrich) were added to each well. If not stated otherwise, growth was monitored in technical triplicates in a BioTek Epoch 2 microtiter plate reader (BioTek, Bad Friedrichshall, Germany) by absorbance measurements (OD600) of each well every ∼10 minutes with intermittent orbital and linear shaking. As previously established empirically for the instrument, blank measurements were subtracted and OD600 measurements were converted to cuvette OD600 values by multiplying with a factor of 4.35. Growth curves represent the average of technical triplicate measurements and were plotted in python with matplotlib using a script available on github. All growth experiments were performed at least in triplicate.

#### Whole-genome sequencing

For whole genome sequencing, the DNeasy Blood and Tissue kit from QIAGEN was used with the equivalent of 1 ml of OD 1 of stationary phase overnight LB culture. Genomic DNA at a concentration no lower than 50 ng/µl was shipped at room temperature to Novogene (illumina) or Plasmidsaurus (nanopore) for sequencing. The resulting data was analyzed with breseq version 0.38.3, bowtie version 2.5.4 and R version 4.5.1, using the -x flag in the case of nanopore data.

### 13C-labeling of proteinogenic amino acids

For the labeling analysis the strains were cultured in 3 ml M9 minimal medium supplemented with 20 mM ^13^C glycerol and 5 mM ^12^C 2-oxoglutarate, methylaspartate, mesaconate, or methylsuccinate. After late exponential phase was reached, an amount equivalent to 1 ml of OD 1 was collected, washed with water three times and resuspended in 1 ml 6 M HCl to hydrolyse the biomass at 95 °C overnight. Then, the samples were completely dried in a thermoblock at 95 °C, resuspended in 100 μl H_2_O and centrifuged to remove insoluble particles.

#### LC-MS

Amino acids were analyzed using HRES LC-MS, performing the chromatographic separation on an Agilent Infinity II 1260 HPLC system with a ZicHILIC SeQuant column (150 × 2.1 mm, 5 µm particle size, 100 Å pore size) connected to a ZicHILIC guard column (20 × 2.1 mm, 5 µm particle size) (Merck KgAA) at a constant flow rate of 0.3 ml/min. Mobile phase A was 0.1 % formic acid in 99:1 water:acetonitrile (Honeywell) and phase B was 0.1 % formic acid in 99:1 acetonitrile:water. The column oven was set to 25 °C, the autosampler was maintained at 4 °C. An injection volume of 1 µl was used. The mobile phase profile consisted of the following steps and linear gradients: 0 to 12 min from 90 to 70% B; 8 12 to 15 min from 70 to 20% B; 15 to 17 min constant at 20% B; 17 to 17.1 min from 20 to 90% B; 17.1 to 20 min constant at 90% B. An Agilent 6550 ion funnel Q-TOF mass spectrometer was used in positive mode with a duel jet stream electrospray ionization source and the following conditions: ESI spray voltage 1000 V, nozzle voltage 900 V, sheath gas 200° C at 9 L/min, nebulizer pressure 35 psi, and drying gas 130° C at 12 L/min. The TOF was calibrated using an ESI-L Low Concentration Tuning Mix (Agilent) before measurement (residuals less than 2 ppm for five reference ions) and was recalibrated during a run using 121.050873 m/z as reference mass. MS data were acquired with a scan range of 50-250 m/z. LC-MS data were analyzed using MassHunter Qualitative Analysis software (Agilent).

#### Expression and purification of HtIDH and HtOGC

N-terminally 6x His-tagged HtIDH, plasmid pTE5501, and N-terminally 10x His-tagged HtOGCL, plasmid pTE5502, were produced from their respective plasmids with *E. coli* BL21-AI as an expression host. N-terminally 10x His-tagged HtOGCS, plasmid pTE5503, was coexpressed with untagged HtBirA, plasmid pTE5504 with *E. coli* BL21-AI as an expression host. The cells were transformed with the respective expression plasmids and then plated on LB agar containing selective antibiotic and grown overnight. The colonies were used to inoculate 8 L of auto induction TB medium. The expression culture was incubated at 25 °C under shaking at 120 rpm for 16 h. The cells were harvested by centrifugation at 5,000 *g* for 10 min at 20 °C. The pellet was optionally stored at –20 °C. The cells were resuspended in a 1:2 ratio (wt/wt) in buffer A (50 mM sodium phosphate, pH 7.9, and 500 mM NaCl) containing DNase I (Merck, Darmstadt, DE) and a Pierce Protease inhibitor tablet (Thermo Scientific, Dreieich, DE). The cells were lysed with a Microfluidizer (LM-10 H10Z, Microfludics, Westwood, USA) at 16,000 PSI for three passes on ice. For HtOGCS coexpresssed with HtBirA, Biotin and Mg_2_-ATP were added dropwise to the crude lysate to yield a final concentration of 1 mM Biotin and 5 mM Mg_2_-ATP and the resulting mixture was then incubated for 30 min at 37°C at 120 rpm. The lysates were clarified by ultracentrifugation, at ∼100,000 *g* for 30 min at 20 °C, and then filtered through a 0.45 μm syringe filter. At room temperature, the clarified lysate was loaded under gravity flow onto a 35 mL protino column, which was packed with ∼10 mL of Ni-NTA (Thermo Scientific) and was preequilibrated with buffer A. Nonspecifically bound proteins were washed away with 50 mL of 10% buffer B (50 mM sodium phosphate, pH 7.9, 500 mM NaCl, and 500 mM imidazole). Proteins were eluted with 100% buffer B. HtIDH was concentrated in Amicon Ultra 15 mL Centrifugal Filters (Merck Darmstadt, DE) to ∼50 mg/mL and then buffer exchanged using a PD-10 that was equilibrated with 50 mM MOPS pH 7.4, 200 mM NaCl, and 5 mM MgCl_2_. Glycerol was added to the protein stock to result in a final concentration of 20% glycerol (v/v). The protein was then flash frozen and stored at -70 °C until further use.

HtOGCS and HtOGCL were concentrated in Amicon Ultra 15 mL Centrifugal Filters (Merck) to 10-15 mg/mL and then buffer exchanged using a PD-10 that was equilibrated with 20 mM Tris-HCl pH 7.9,150 mM NaCl, and 5 mM MgCl_2_,. HtOGCS (∼20 mg) was then mixed with HtOGCL (∼30 mg) and incubated at 37 °C at 120 rpm for 30 min. The mixture was then loaded onto a preequilibrated HiLoad 16/60 200 pg Superdex (GE Life Science) column (20 mM Tris-HCl pH 7.9, 150 mM NaCl, and 5 mM MgCl_2_) at room temperature. Fractions corresponding to the protein complex eluted at ∼50 mL. The composition of the eluted fractions were analyzed by SDS-PAGE. Fractions containing ∼1:1 ratio of OGCS: OGCL were combined. Concentrating the enzyme beyond ∼2 mg/mL resulted in precipitation and was therefore avoided. Glycerol was added to the protein stock to result in a final concentration of 20% glycerol (v/v). The resulting HtOGC complex was then flash frozen and stored at -70 C until further use. All protein concentrations were determined by Bradford Assay using the Quick Start Bradford Protein Assay kit (Bio-Rad Laboratories, Hercules, USA).

#### HtOGC and HtIDH coupled assays

The assay temperature was set to 60 °C. The specific activity for HtOGC was measured in a coupled assay with HtIDH. Assays contained 100 mM Tris pH 8, 0.3 mM NADH, 50 mM NaHCO_3_, 25mM MgCl_2_, 5mM ATP, 100 ug/mL HtOGC, 25 ug/mL HtIDH, and varying concentrations of 2-oxoglutarate, in a final volume of 0.15 mL. The assays were incubated at 60 °C for 2 minutes before HtIDH was added and the measurement was started. After ∼30 s Mg-ATP was added to assay. The oxidation of NADH (ε_NAD(P)H,340nm_ = 6.22 mM^−1^ cm^−1^) was followed at 340 nm on a Cary 60 UV–Vis photospectrometer (Agilent, Santa Clara, USA) in quartz cuvettes with a pathlength of 10 mm (Hellma Optik GmbH, Jena, Germany). Measurements were performed in triplicates and V_max_ and K_m_ were calculated using Beer-Lambert’s Law. Data was fit using Michaelis-Menten function in Prism 8 (Graphpad, San Diego, USA).

#### HtIDH Isocitrate decarboxylation assays

The assay temperature was set to 37 °C. Assays contained 100 mM Tris pH 8, 4 mM NAD(P)^+^, 200 μg/mL or 400 μg/mL HtIDH for NADP^+^ and 20 μg/mL or 40 μg/mL HtIDH for NAD^+^ in a final volume of 0.2 mL. The assays were incubated at 37 °C for 1 min and then HtIDH was added and the measurement was initiated. After ∼30 s DL-isocitric acid was added to assay. The reduction of NAD(P)^+^ (ε_NAD(P)H,340nm_ = 6.22 mM^−1^ cm^−1^) was followed at 340 nm on a Cary 60 UV–Vis photospectrometer (Agilent, Santa Clara, USA) in quartz cuvettes with a pathlength of 10 mm (Hellma Optik GmbH, Jena, Germany).

Measurements were performed in triplicates and V_max_ and K_m_ were calculated using Beer-Lambert’s Law. Data was fit using Michaelis-Menten function in Prism 8 (Graphpad, San Diego, USA).

#### HtIDH 2-oxoglutarate carboxylation assays

The assay temperature was set to 70 °C. Assays contained 100 mM Tris pH 8, 0.3 mM NADH, 50 mM NaHCO_3_, and 2.75 mg/mL HtIDH in a final volume of 0.2 mL. The assays were incubated at 70 °C for 2 min and then HtIDH was added and the measurement was initiated. After ∼30 s 2-oxoglutarate was added to assay. The oxidation of NADH (ε_NAD(P)H,340nm_ = 6.22 mM^−1^ cm^−1^) was followed at 340 nm on a Cary 60 UV–Vis photospectrometer (Agilent, Santa Clara, USA) in quartz cuvettes with a pathlength of 10 mm (Hellma Optik GmbH, Jena, Germany). Measurements were performed in triplicates and V_max_ and K_m_ were calculated using Beer-Lambert’s Law. Data was fit using Michaelis-Menten function in Prism 8 (Graphpad, San Diego, USA).

## Acknowledgements

The authors would like to thank Anton Bunschoten for granting access to the Bionanotechnology lab facilities, Merlin Cotessat for assistance with organic synthesis and NMR, Bob Rose and Amy Grunden for providing the pQE1-OGC plasmid, Shanshan Luo for providing the pTE3288 and pTE3236 plamids and for assistance with the sdhA in vitro assay, and Tim Althuis for valuable input on the figures. VR is funded by the VLAG PhD school of Wageningen University, NJC acknowledges funding from the ERC StG FASTFIX by the European Research Council.

## Author contributions

VR conceptualized the project, designed and analyzed all cycles, performed most experiments, analyzed the data, and wrote the manuscript. HSM genetically engineered strain V1 and analyzed the labeling data. JBS performed the in vitro OGC and IDH assays. SQS cloned and tested all glutamate mutase plasmids, NP performed the LC-MS measurement of amino acids. TJE, SDA, and NJC supervised the research and acquired funding. All authors contributed to editing the manuscript.

## Conflict of interest statement

NJC is an advisor to the start-up companies Farmless and Novya Biotech, but these companies were not in any way involved in this study. The other authors declare no conflict of interest.

---

**Figure S1.**
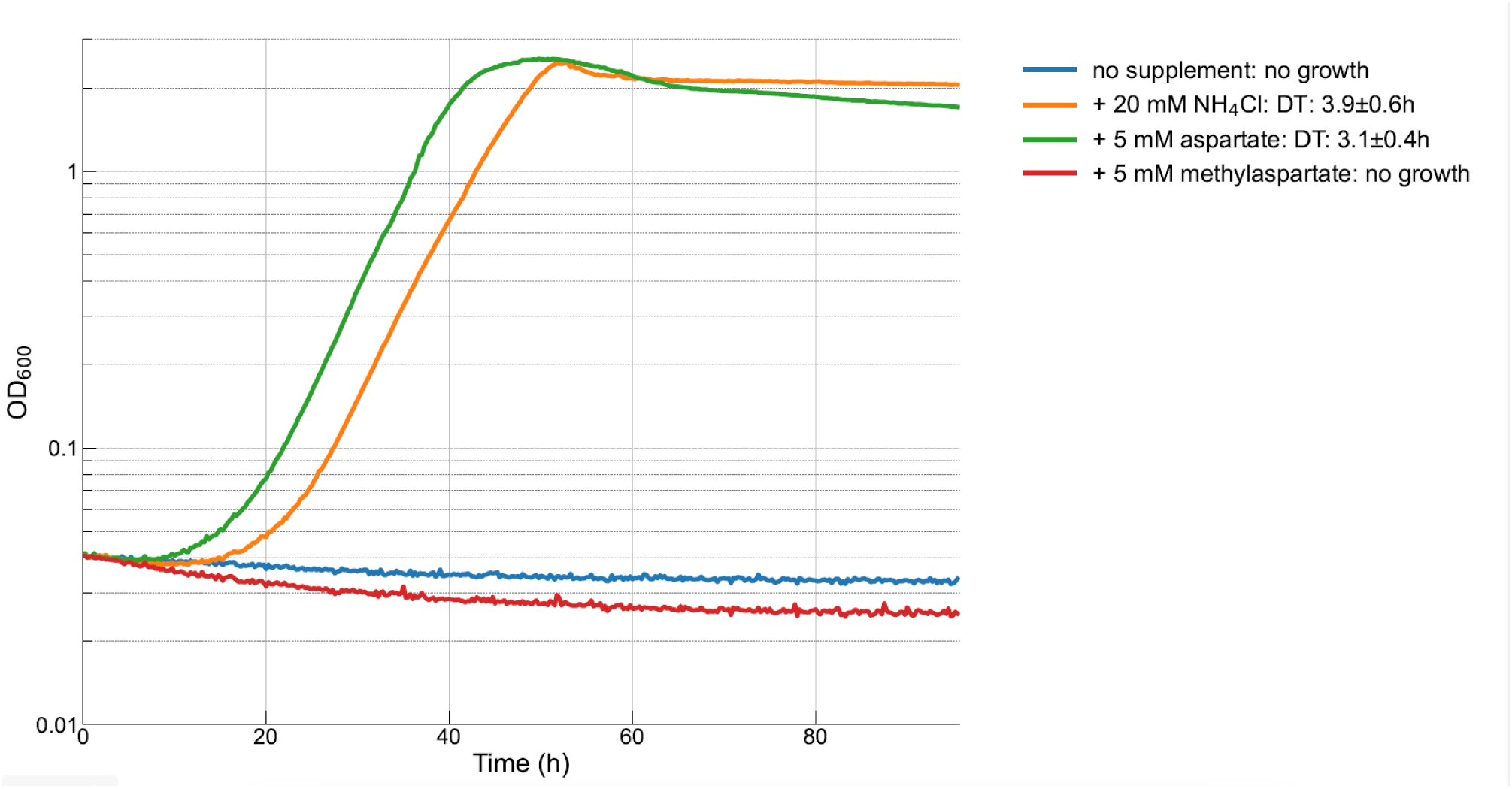
Growth of *E. coli* overexpressing the native aspartate ammonia lyase (aspA) in M9 devoid of a nitrogen source. Aspartate supplementation releases ammonium which can be used for growth, while methylaspartate does not, indicating that the latter is not a substrate for this enzyme. All growth curves are the average of at least three replicates.

**Figure S2.**
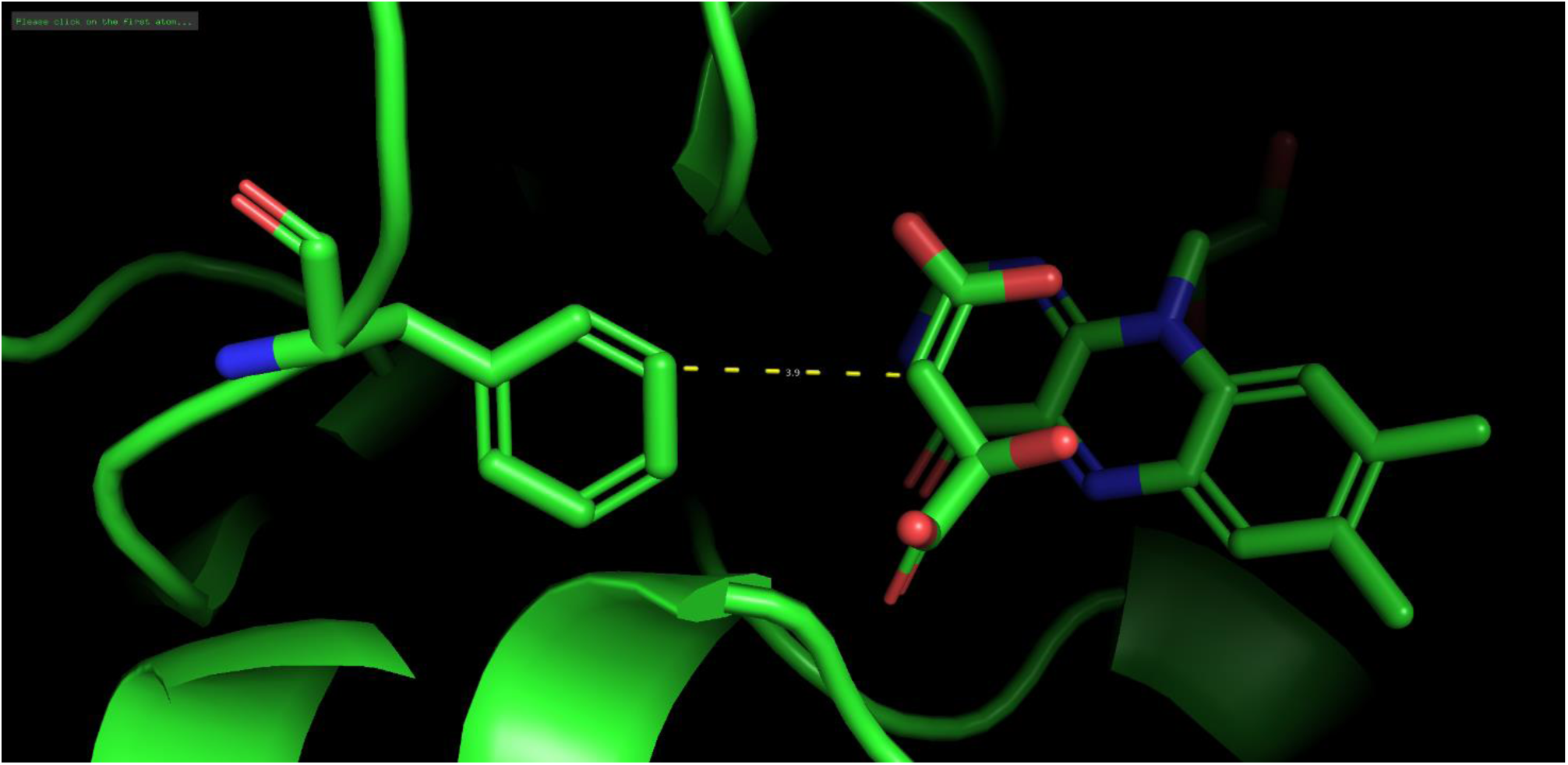
*E. coli* SdhA (PDB: 2wdv) showing the distance between F119 and the substrate analog (Z,2R)-2,4-dihydroxy-4-oxido-but-3-enoate (TEO), a malate-like molecule that cannot act as a substrate for the enzyme. A distance of 3.9 Å likely indicates a van der Waals interaction. The extra methyl group in methylsuccinate would likely interfere with this interaction.

**Figure S3.**
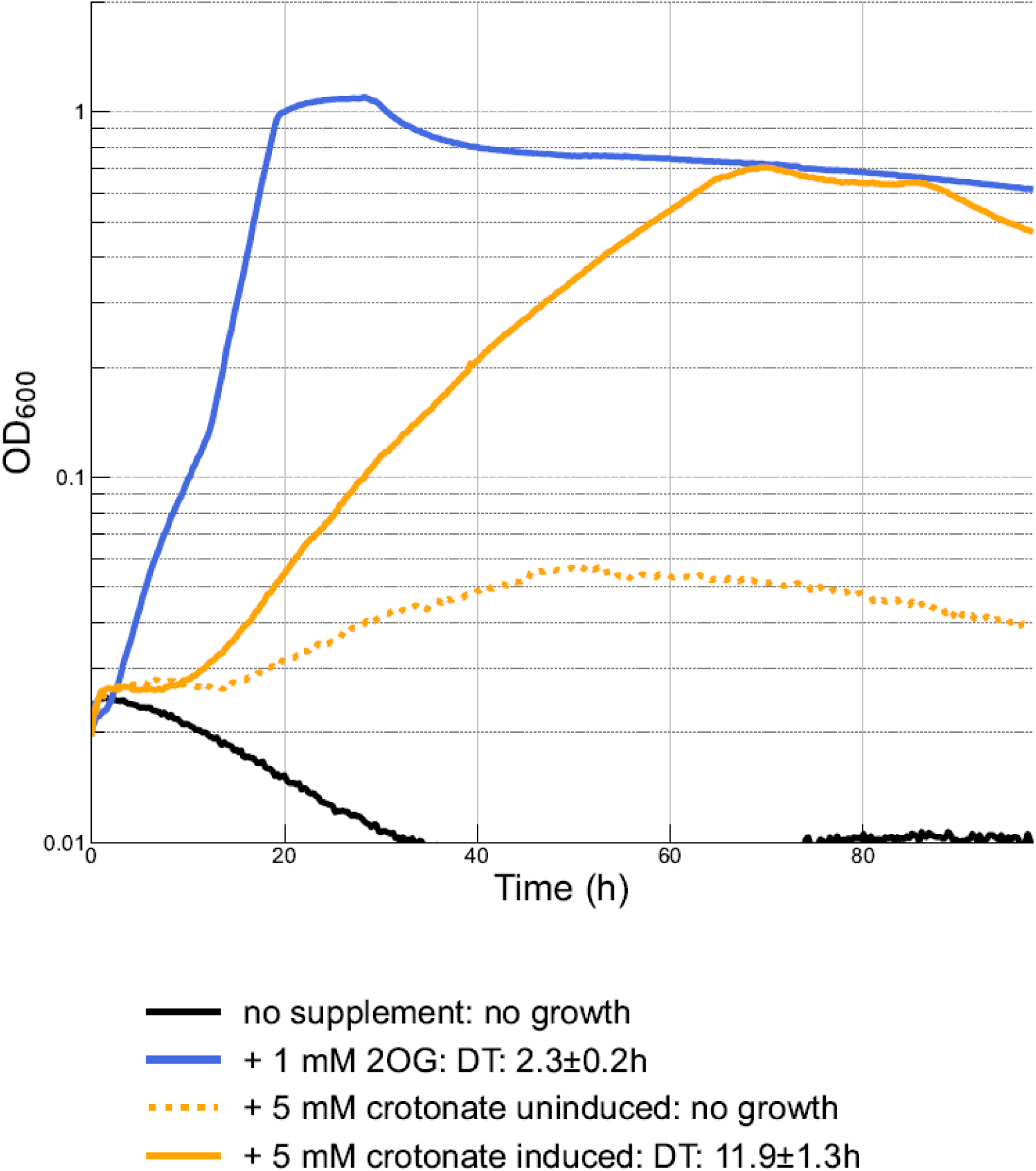
Growth of the V2 strain in M9N100 with 10 mM gluconate, 10 µM B_12_, 50 µM IPTG, 5% CO_2_ and supplemental carbon sources as indicated in the legend. Dotted lines indicate that IPTG was omitted as negative control. 2OG, 2-oxoglutarate; DT, doubling time.

**Figure S4.**
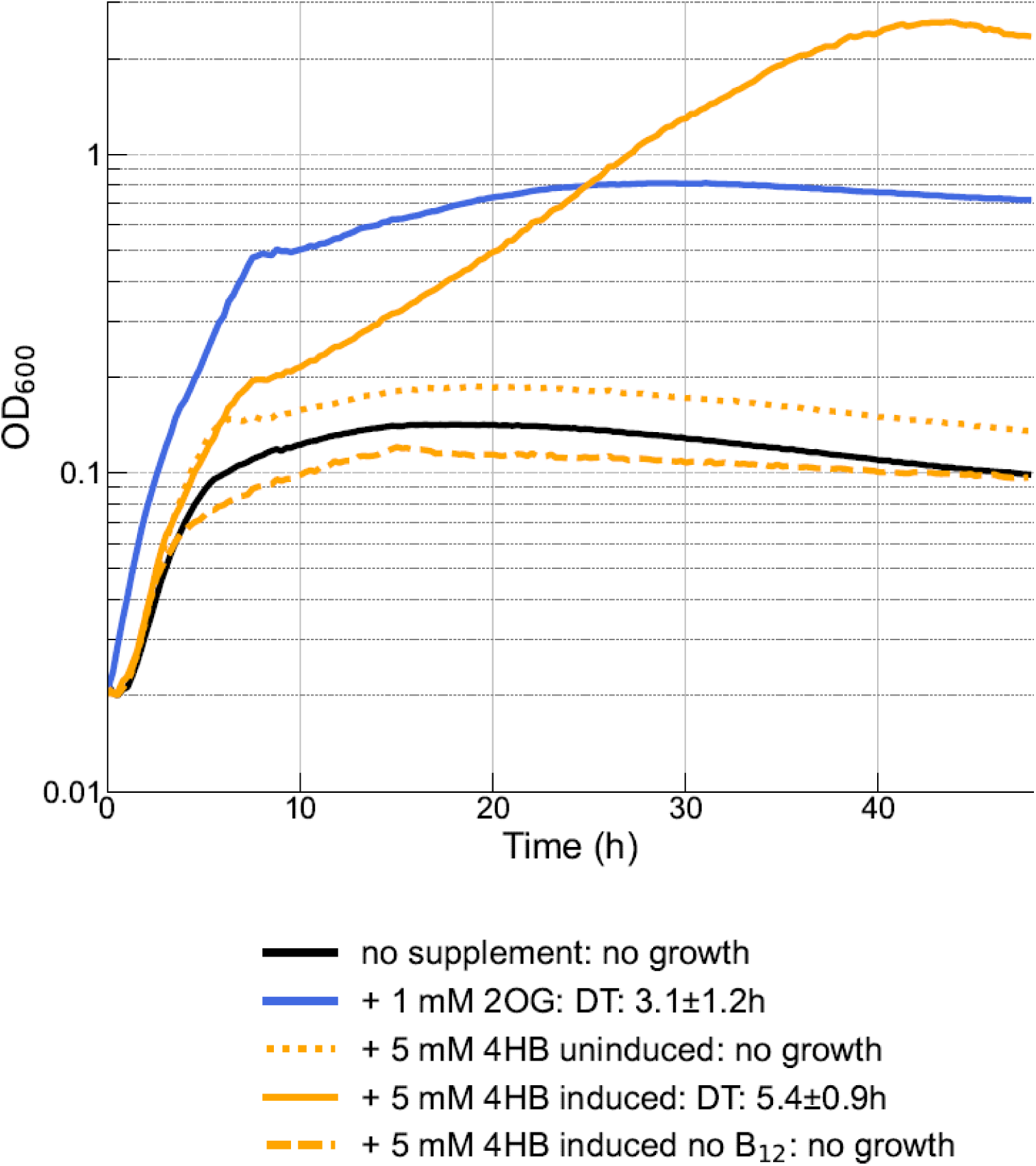
Growth of the V3 strain carrying pTE3236E in M9N100 with 10 mM gluconate, 10 µM B_12_, and 50 µM IPTG, and supplementary carbon sources as indicated in the legend at 5% CO_2_ concentration. Dotted and dashed lines indicate the omission of IPTG or B_12_ from the medium, respectively. 2OG, 2-oxoglutarate; 4HB, 4-hydroxybutyrate; DT, doubling time.

**Figure S5.**
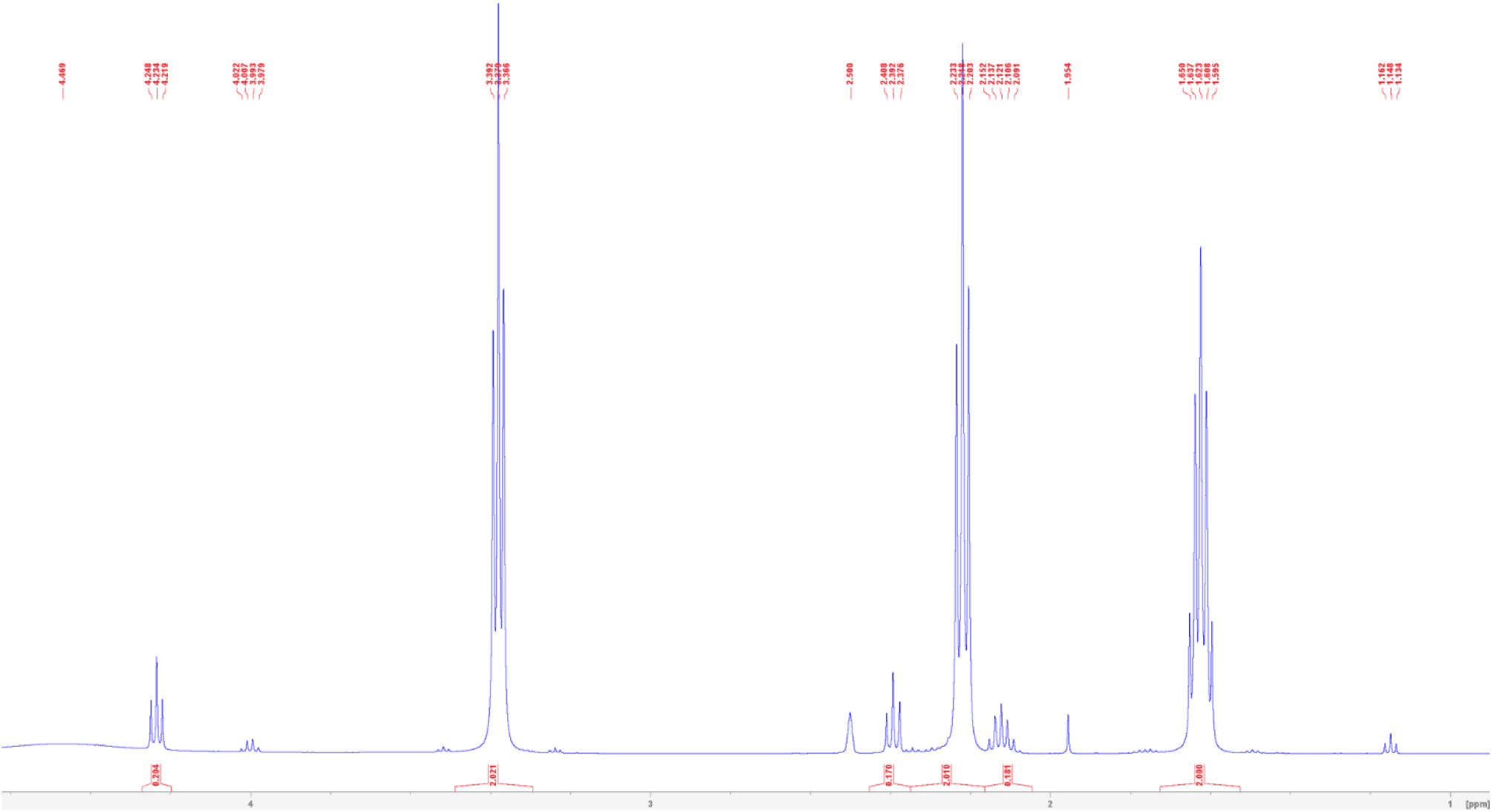
NMR spectrum of 4-hydroxybutyrate in DMSO after deesterification of γ-butyrolactone, extraction in ethyl acetate, and solvent evaporation.

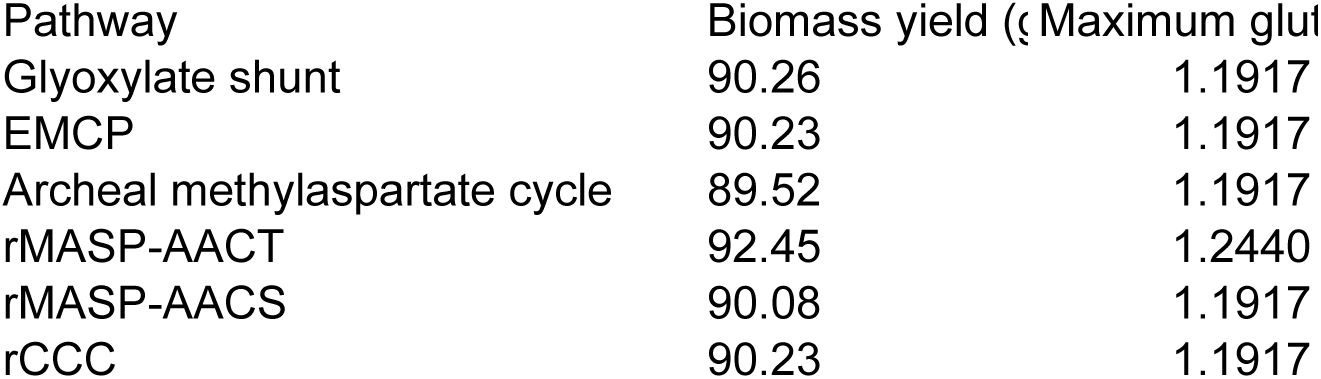

| Biomass Yield (gCDW/mol acetate) | Maximum glutamate yield (mol/mol acetate) |
| --- | --- |
| 19.57 | 0.2958 |
| 19.67 | 0.2973 |
| 18.71 | 0.28 |
| 21.07 | 0.3235 |
| 19.54 | 0.2958 |
| 18.99 | 0.2683 |

## Notes

https://git.wur.nl/vittorio.rainaldi/r_masp

### Competing Interest Statement

The authors have declared no competing interest.

