## Supplementary Data Files for "Modular in vivo engineering of the reductive methylaspartate cycles for synthetic CO_2_ fixation": Manuscript.docx

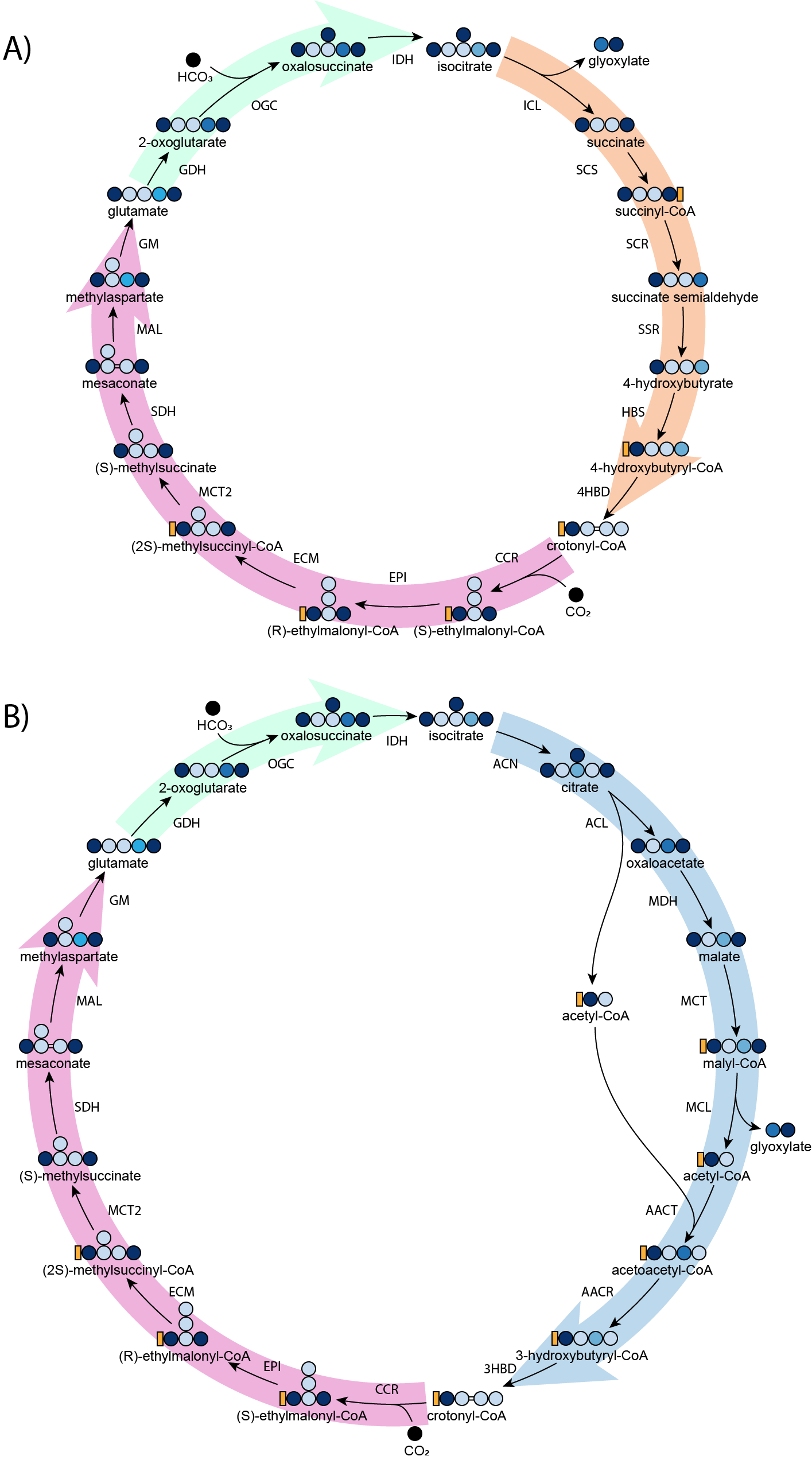

**Figure 1**. The two different pathway architectures of the reductive methylaspartate cycle. (A) via isocitrate lyase (ICL) and (B) via acetoacetyl-CoA thiolase (AACT). 3HBD, 3-hydroxybutyryl-CoA dehydratase; 4HBD, 4-hydroxybutyryl-CoA dehydratase; ACL, ATP citrate lyase; ACN, aconitase; AACR, acetoacetyl-CoA reductase; AACT, acetoacetyl-CoA thiolase; CCR, crotonyl-CoA carboxylase; ECM, ethylmalonyl-CoA mutase; EPI, ethylmalonyl-CoA epimerase; ICL, isocitrate lyase; IDH, isocitrate dehydrogenase; GDH, glutamate dehydrogenase; GM, glutamate mutase; HBS, 4-hydroxybutyryl-CoA synthase; HBD, 4-hydroxybutyryl-CoA dehydratase. MAL, mesaconate ammonia lyase; MCL, malyl-CoA lyase; MCT, malate CoA transferase; MCT2, mesaconyl-CoA transferase; MDH, malate dehydrogenase; SDH, methylsuccinate dehydrogenase; OGC, 2-oxoglutarate carboxylase; SCS, succinyl-CoA synthetase; SCR, succinyl-CoA reductase; SSR, succinate semialdehyde reductase.

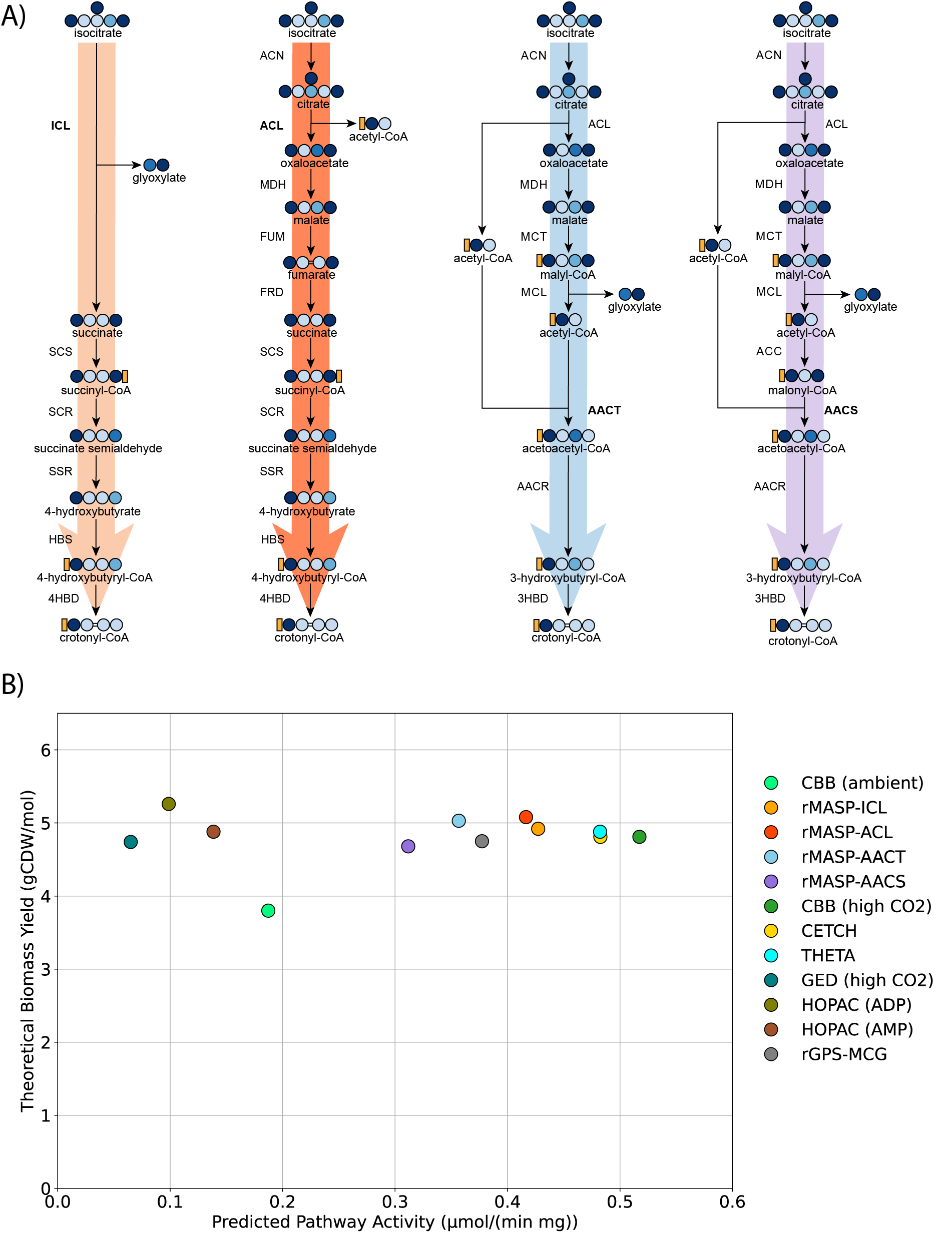

**Figure 2.** (A) Four modules to regenerate crotonyl-CoA from isocitrate (B) Computational prediction of biomass yield (calculated using FBA) and pathway activity (calculated using ECM) for each variant. Yields are calculated per mol NADH. The ambient CO_2_ condition for the CBB assumes a 20% rate of rubisco oxygenation^42^. AACS, acetoacetyl-CoA synthase; ACC, acetyl-CoA carboxylase; FRD, fumarate reductase; other enzyme abbreviations are the same as in figure 1.

**In vivo implementation of the glutamate module**

The glutamate module is shared across all cycle variants and was never demonstrated in vitro or in vivo (Figure 1, pink arrows). For this reason, we used it as the starting point for in vivo implementation in a glutamate auxotroph strain.

In this work, we used an *E. coli* isocitrate dehydrogenase *(*Δ*icd*) knockout strain from the KEIO collection^43^. This strain is unable to synthesize 2-oxoglutarate (2OG) and hence is auxotrophic for glutamate and glutamate-derived amino acids. This strain can only grow on glycerol minimal medium with 2OG or glutamate supplementation (figure 3A). Using this strain, we tested the module in a stepwise manner, starting from the conversion of methylaspartate to glutamate, a reaction catalyzed by the B_12_-dependent enzyme glutamate mutase.

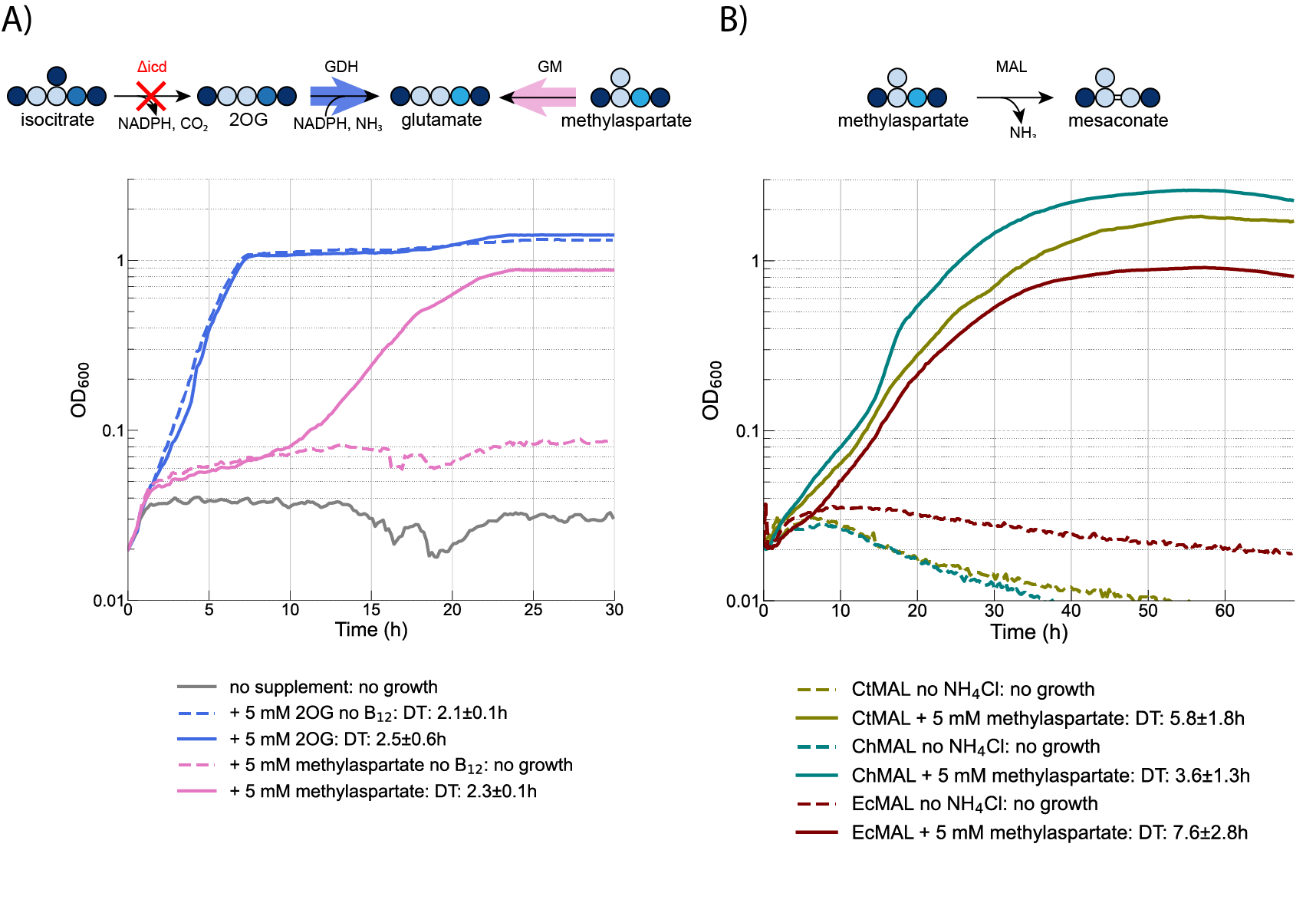

**Figure 3.** (A) Growth of an *E. coli* glutamate auxotroph (Δ*icd*) via B_12_-dependent glutamate mutase *sanUV* from *S. ansochromogenes*. The strain requires both methylaspartate and B_12_ to produce glutamate. The strain was grown in M9 minimal medium with 20 mM glycerol as a main carbon source and supplements as indicated in the legend. All experiments were performed under fully aerobic conditions, demonstrating that the chosen glutamate mutase is not inactivated by oxygen. (B) In vivo demonstration of functional mesaconate ammonia lyase (MAL) expression. The strains were grown in M9 minimal medium devoid of a nitrogen source (M9N0). Supplementation with methylaspartate allows growth via deamination to mesaconate. Growth curves are the average of at least three technical replicates. DT, doubling time; GDH, glutamate dehydrogenase; GM, glutamate mutase; MAL, methylaspartate ammonia lyase.

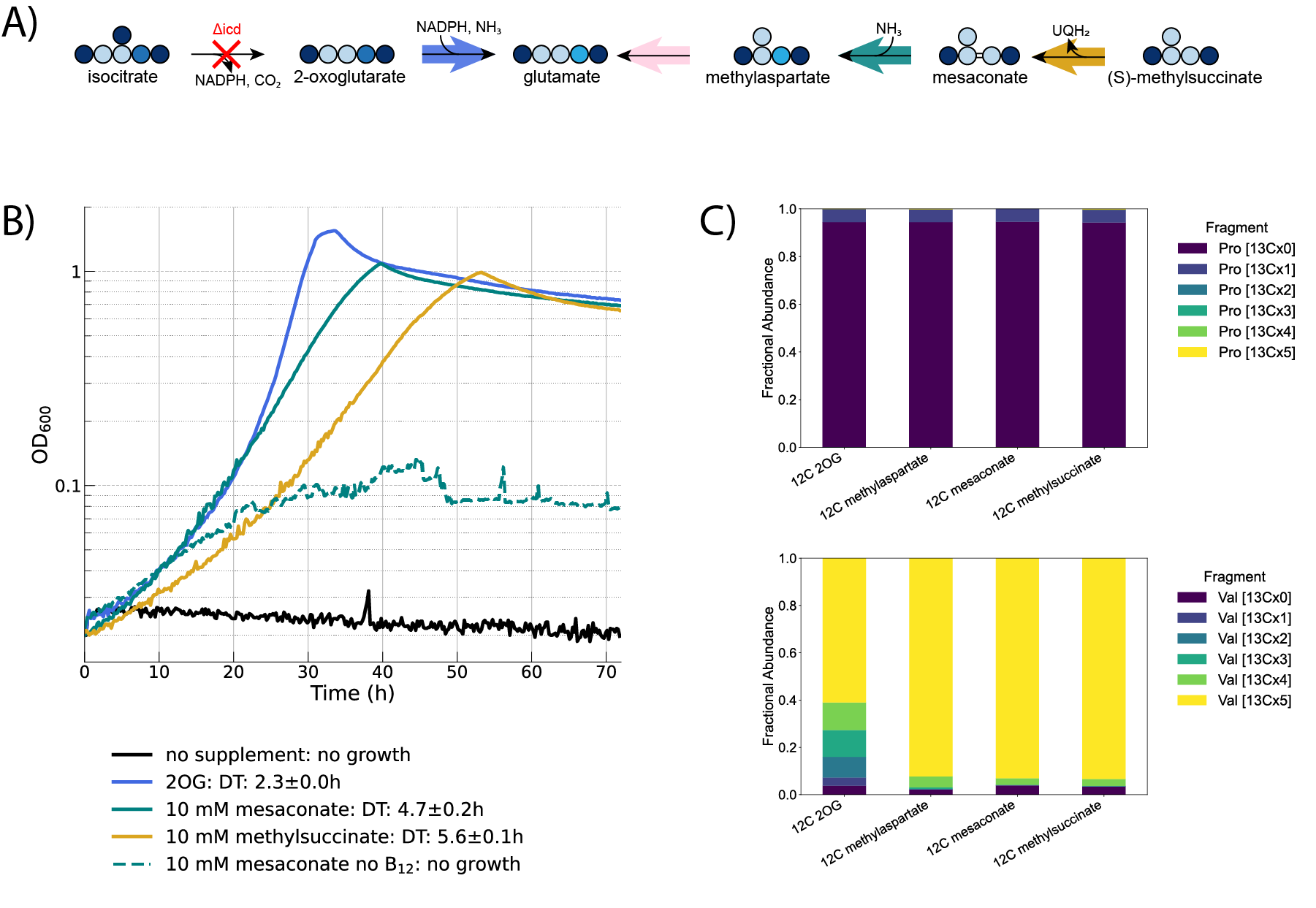

**Figure 4.** (A) Metabolic scheme of the growth of an *E. coli* glutamate auxotroph (Δ*icd*) via a B_12_-dependent glutamate mutase. The strain cannot grow using glycerol as a sole carbon source, and requires supplementation of a source of glutamate. (B) Growth curves when different supplements are added. Growth is B_12_-dependent, as indicated by the no growth phenotype when B_12_ is absent from the medium (dashed grey line) (C) Isotopic labeling data showing the incorporation of unlabeled carbon sources into biomass. All strains were grown in M9 with 20 mM fully ^13^C labeled glycerol and unlabeled (^12^C) supplementary carbon sources as indicated in the figure. Proline is derived from glutamate and hence expected to be fully unlabeled, whereas valine is derived from pyruvate and expected to be fully labeled. The 2OG condition shows that this metabolite is partially converted to pyruvate via the TCA cycle.

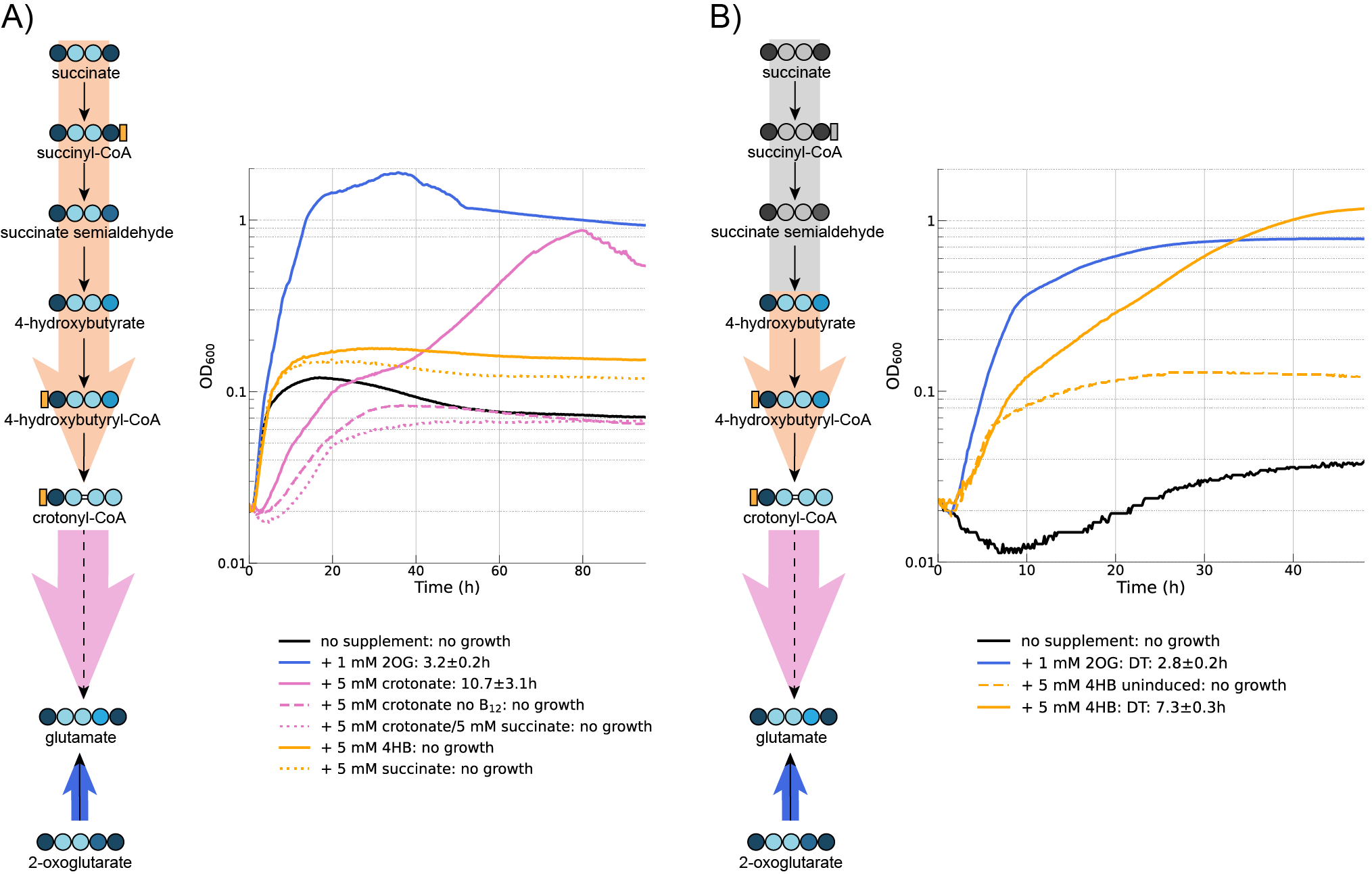

**Figure 5. (A)** Crotonate-dependent growth of the V3 strain carrying pTE3236C via the full mutase module (solid pink line). Addition of succinate (dotted pink line) inhibits growth, potentially indicating a toxicity effect due to the competition between succinate and methylsuccinate for Sdh activity. Neither succinate nor 4HB (orange lines) support growth in the absence of crotonate **(B)** 4HB-dependent growth of the V3 strain carrying pTE3236D, in which the greyed out enzymes were removed. Experiments were performed in M9N100 with 10 mM gluconate, 10 µM B_12_, 50 µM IPTG, 5% CO_2_ and supplemental carbon sources as indicated in the legend. B_12_ or IPTG were omitted as negative controls, indicated by dashed lines. 2OG, 2-oxoglutarate; 4HB, 4-hydroxybutyrate; DT, doubling time

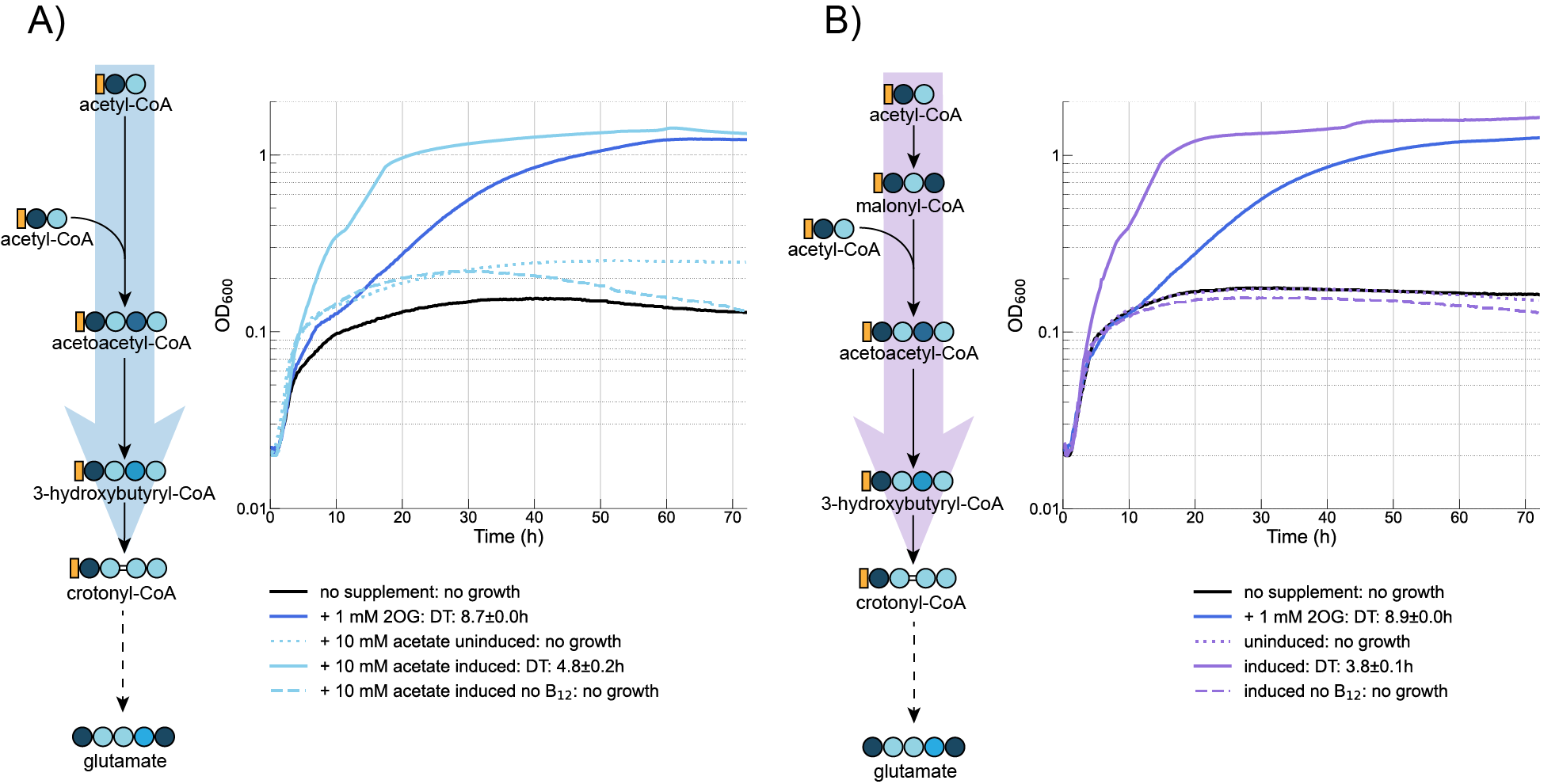

**Figure 6.** Growth of the V3 strain via the AACT (left) and AACS (right) modules for acetate assimilation. Strains were grown in M9N100 with 10 mM gluconate as a main carbon source, 10 µM B_12_, 10 µM cuminic acid, 5% CO_2_ and supplements as indicated in the legend. Dashed or dotted lines indicate the omission of B_12_ or cuminic acid, respectively.

**The CCR from *Methylorubrum extorquens* allows growth at ambient CO_2_ concentration**

Throughout this work, we used the CCR from *R. sphaeroides* (RsCCR) in all our constructs, since this variant was previously used in vitro test and in vivo attempts of the implementation of the THETA cycle. This enzyme is the best characterized CCR, but it shows relatively poor kinetic parameters with regards to affinity for both NADPH and CO_2_ (0.7 mM and 0.2 mM, respectively). The concentration of NADPH in *E. coli* is known to be around 0.1 mM^57^ and the concentration of dissolved CO_2_ at atmospheric pressure is between 10 and 20 µM^58^, meaning that the enzyme is likely operating well below its maximal velocity under ambient CO_2_ conditions. We reasoned that this likely resulted in the observed lack of growth observed without high CO_2_ partial pressure or bicarbonate supplementation. Since functionality at atmospheric CO_2_ concentrations is a core criterion for synthetic CO_2_ fixation in photosynthetic conditions with ambient CO_2_ and O_2_, we decided to test an alternative CCR from *Methylorubrum extorquens* (MeCCR). While the affinity for CO_2_ of this specific isozyme has not been reported, we expected better performance, since *M. extorquens* grows via the EMCP using the serine cycle in an environment (the phyllosphere) where CO_2_ concentration is not higher than normal, and in fact could even be CO_2_ depleted due to the photosynthetic activity of leaves. By substituting RsCCR with MeCCR we observed 4HB-dependent growth in ambient cultivation conditions (Fig. 7). Notably, the doubling time improved from 7.3 h to 5.7 h, indicating that MeCCR is kinetically faster at ambient CO_2_ than RsCCT at high CO_2_. Furthermore, there is no difference in doubling time between MeCCR at ambient CO_2_ vs high CO_2_ (5.7 h vs 5.4 h, fig. S5) which indicates that ambient CO_2_ is likely saturating.

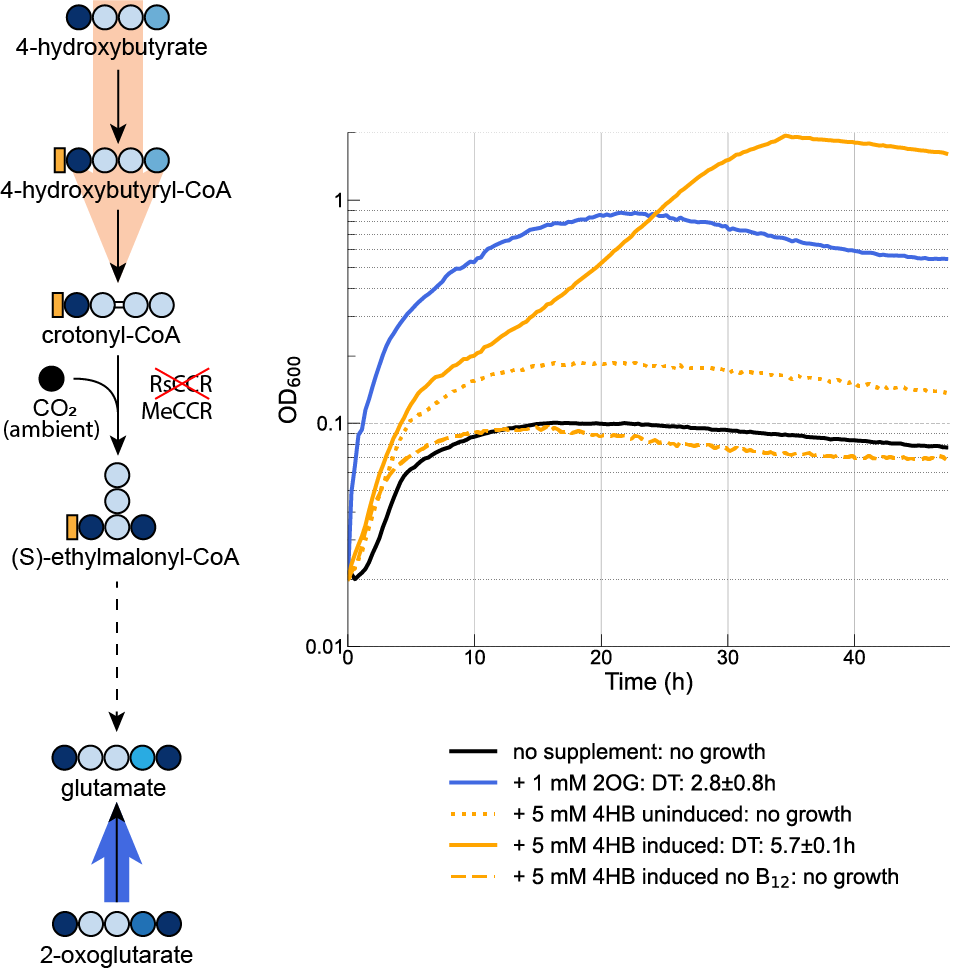

**Figure 7.** Growth of the V3 strain carrying pTE3236E in M9N100 with 10 mM gluconate, 10 µM B_12_, and 50 µM IPTG, and supplementary carbon sources as indicated in the legend under ambient CO_2_ concentration. Dotted and dashed lines indicate the omission of IPTG or B_12_ from the medium, respectively. 2OG, 2-oxoglutarate; 4HB, 4-hydroxybutyrate; DT, doubling time.

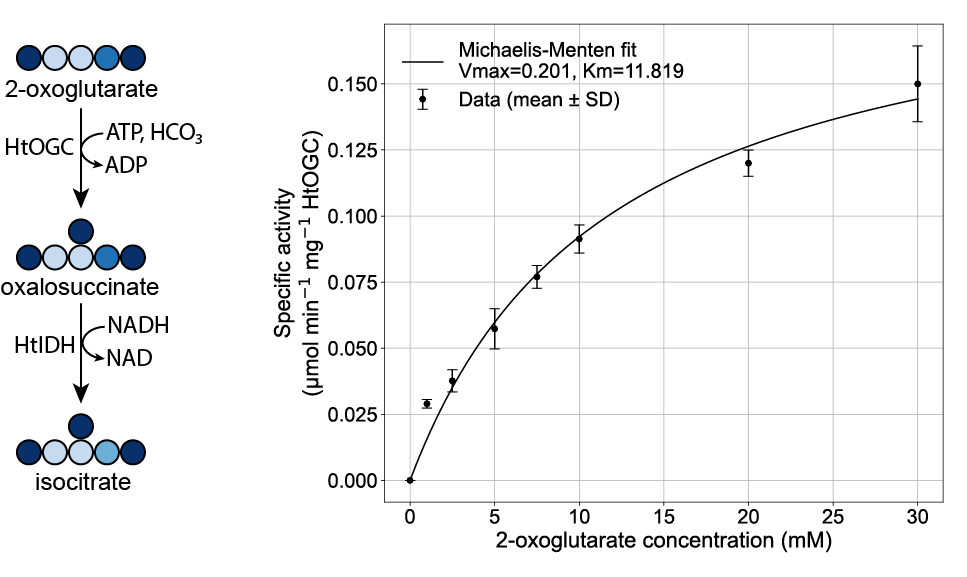

**Figure 8**. In vitro coupled assay of HtOGC and HtIDH after heterologous expression in *E. coli* and purification. The assay was performed at 60 °C in triplicate.

**Discussion**

In this study, we presented the novel rMASP CO_2_ fixation pathways, which have a higher theoretical biomass yield than other natural and synthetic counterparts, such as the CBB, CETCH, THETA and rGPS-MCG cycles. Our design represents a new, previously unexplored family of carbon fixation cycles based on an uncommon carboxylase activity found in hyperthermophilic chemolithotrophs. Until recently, none of the proposed synthetic pathways used this enzyme, showing that new carboxylation reactions can unlock entirely new modes of carbon fixation with the potential to be more efficient than current alternatives. Notably, a comprehensive computational analysis of all possible carbon fixation pathway based on known biochemical reactions recently failed to identify the rMASP cycle^17^. This can be attributed to the fact that the promiscuous activity of succinate dehydrogenase on methylsuccinate is not annotated in databases and hence was not included in that work. This highlights the need for curation of reaction sets based on biochemical expertise when performing such computational analyses.

Pathway activity = $\frac{\mu mol}{\min*mg}$

flux = x $\frac{mM}{s}=1000x\frac{\mu M}{s}=60000x\frac{\mu M}{min}=60000x \frac{\mu mol}{\min*L}$

Enzyme Cost (EC) = y $\frac{g}{L}=1000y\frac{mg}{L}$

Pathway activity = $60000x \frac{\mu mol}{\min*L} \left( \frac{1}{1000y} \right)\frac{L}{mg}=60\frac{x}{y}\frac{\mu mol}{\min*mg} = 60\frac{flux}{EC}$

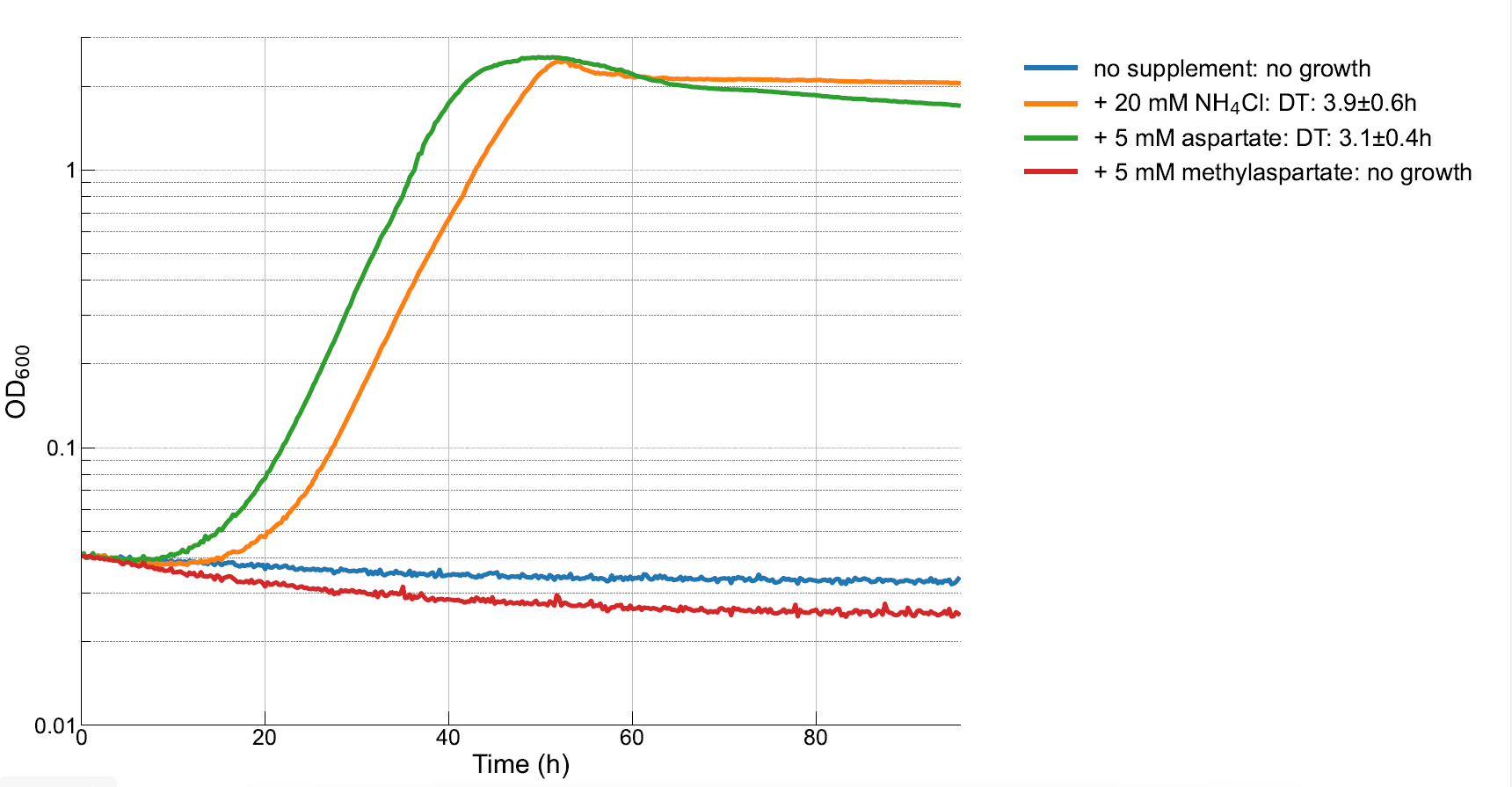

Figure S1. Growth of *E. coli* overexpressing the native aspartate ammonia lyase (aspA) in M9 devoid of a nitrogen source. Aspartate supplementation releases ammonium which can be used for growth, while methylaspartate does not, indicating that the latter is not a substrate for this enzyme. All growth curves are the average of at least three replicates.

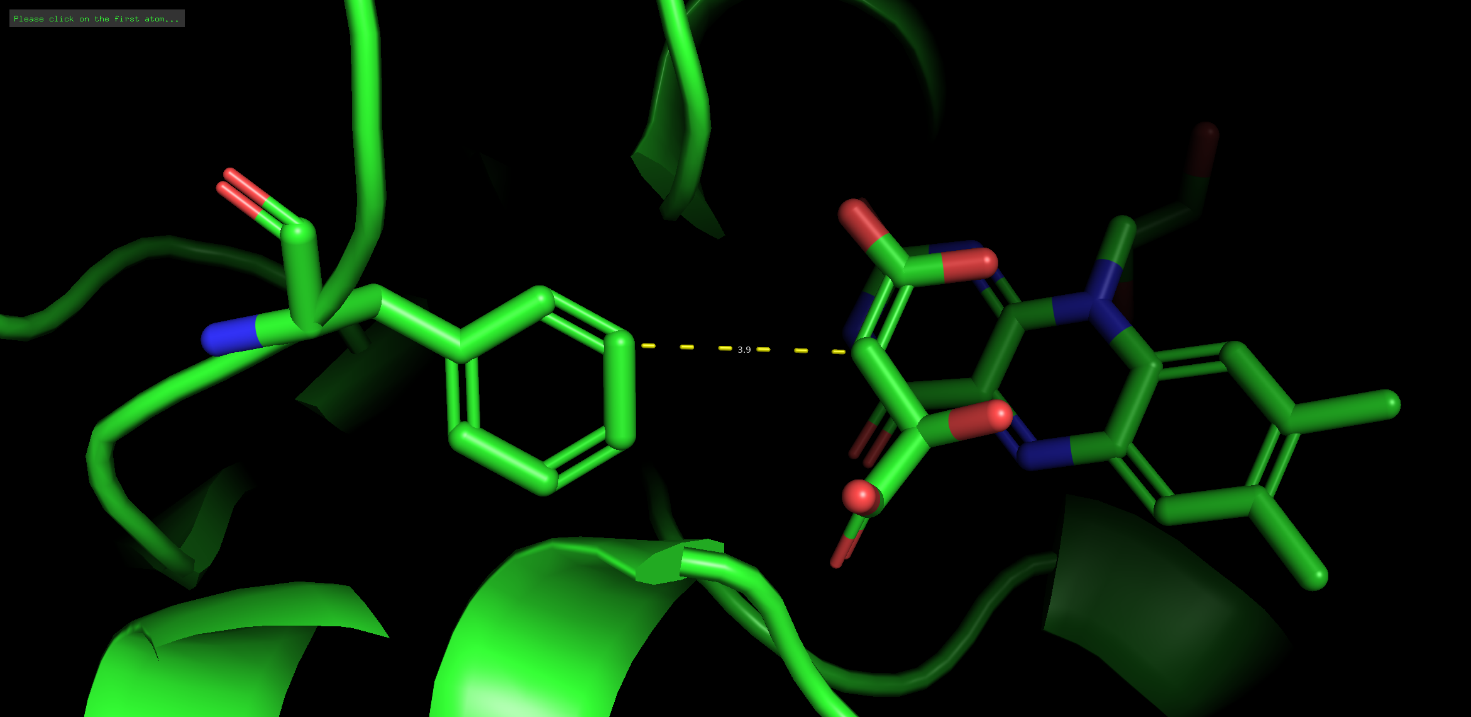

Figure S2. *E. coli* SdhA (PDB: 2wdv) showing the distance between F119 and the substrate analog (Z,2R)-2,4-dihydroxy-4-oxido-but-3-enoate (TEO), a malate-like molecule that cannot act as a substrate for the enzyme. A distance of 3.9 Å likely indicates a van der Waals interaction. The extra methyl group in methylsuccinate would likely interfere with this interaction.

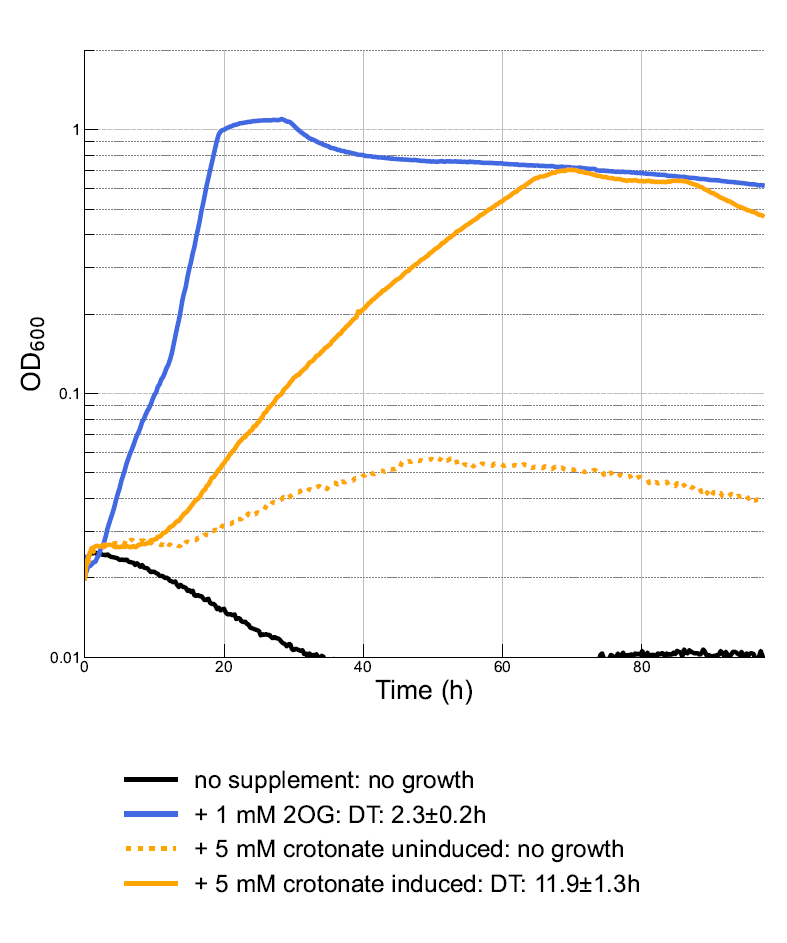

Figure S3. Growth of the V2 strain in M9N100 with 10 mM gluconate, 10 µM B_12_, 50 µM IPTG, 5% CO_2_ and supplemental carbon sources as indicated in the legend. Dotted lines indicate that IPTG was omitted as negative control. 2OG, 2-oxoglutarate; DT, doubling time.

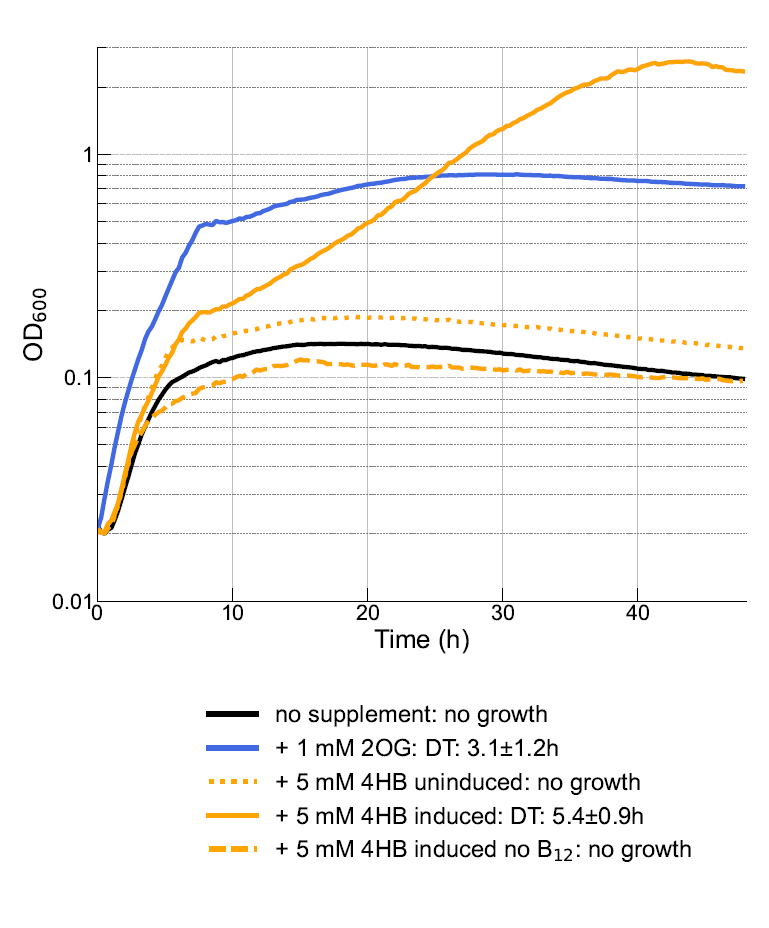

Figure S4. Growth of the V3 strain carrying pTE3236E in M9N100 with 10 mM gluconate, 10 µM B_12_, and 50 µM IPTG, and supplementary carbon sources as indicated in the legend at 5% CO_2_ concentration. Dotted and dashed lines indicate the omission of IPTG or B_12_ from the medium, respectively. 2OG, 2-oxoglutarate; 4HB, 4-hydroxybutyrate; DT, doubling time.

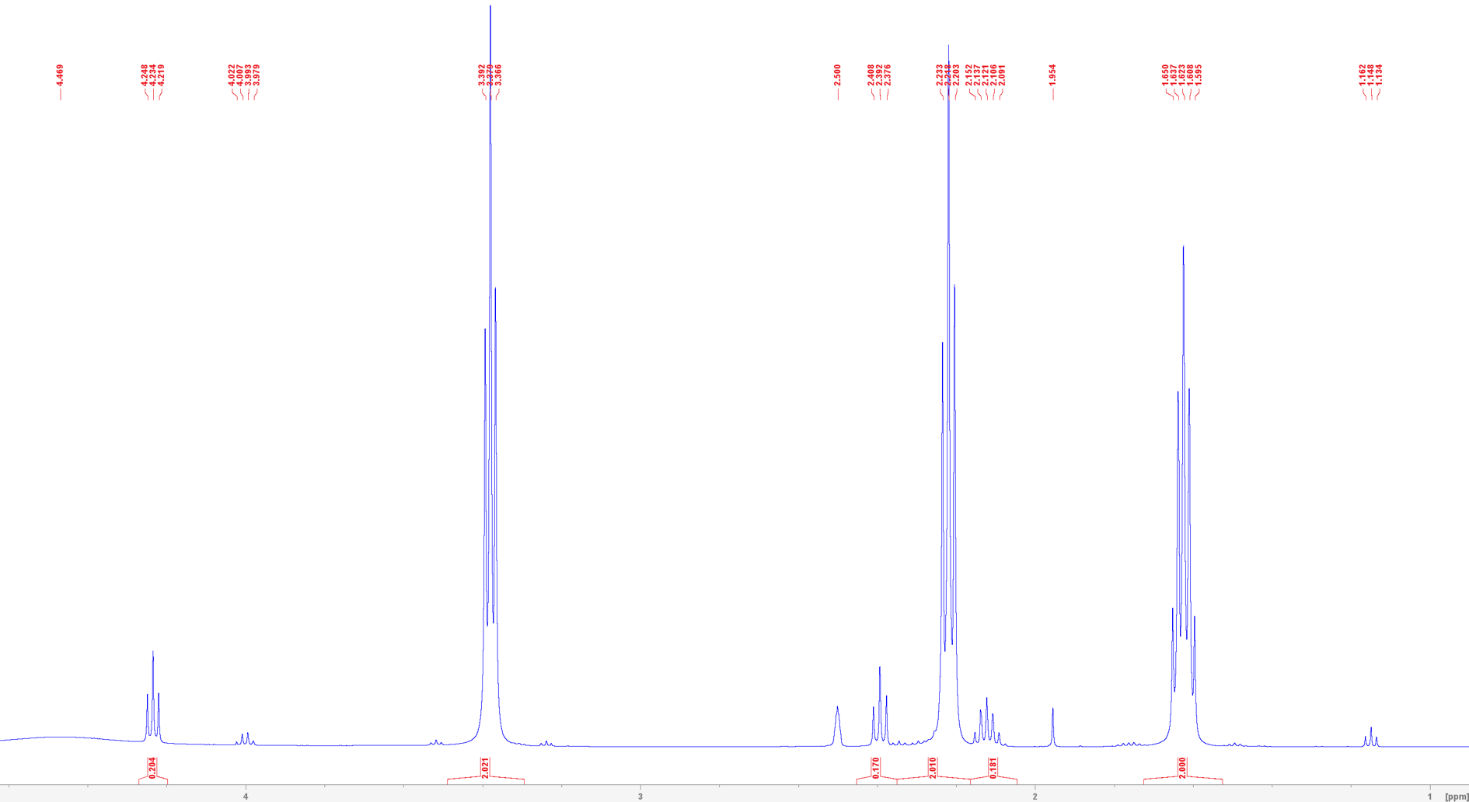

Figure S5. NMR spectrum of 4-hydroxybutyrate in DMSO after deesterification of γ-butyrolactone, extraction in ethyl acetate, and solvent evaporation.
