## Supplementary figures and images for "Modular in vivo engineering of the reductive methylaspartate cycles for synthetic CO_2_ fixation"

### Figure_1.pdf

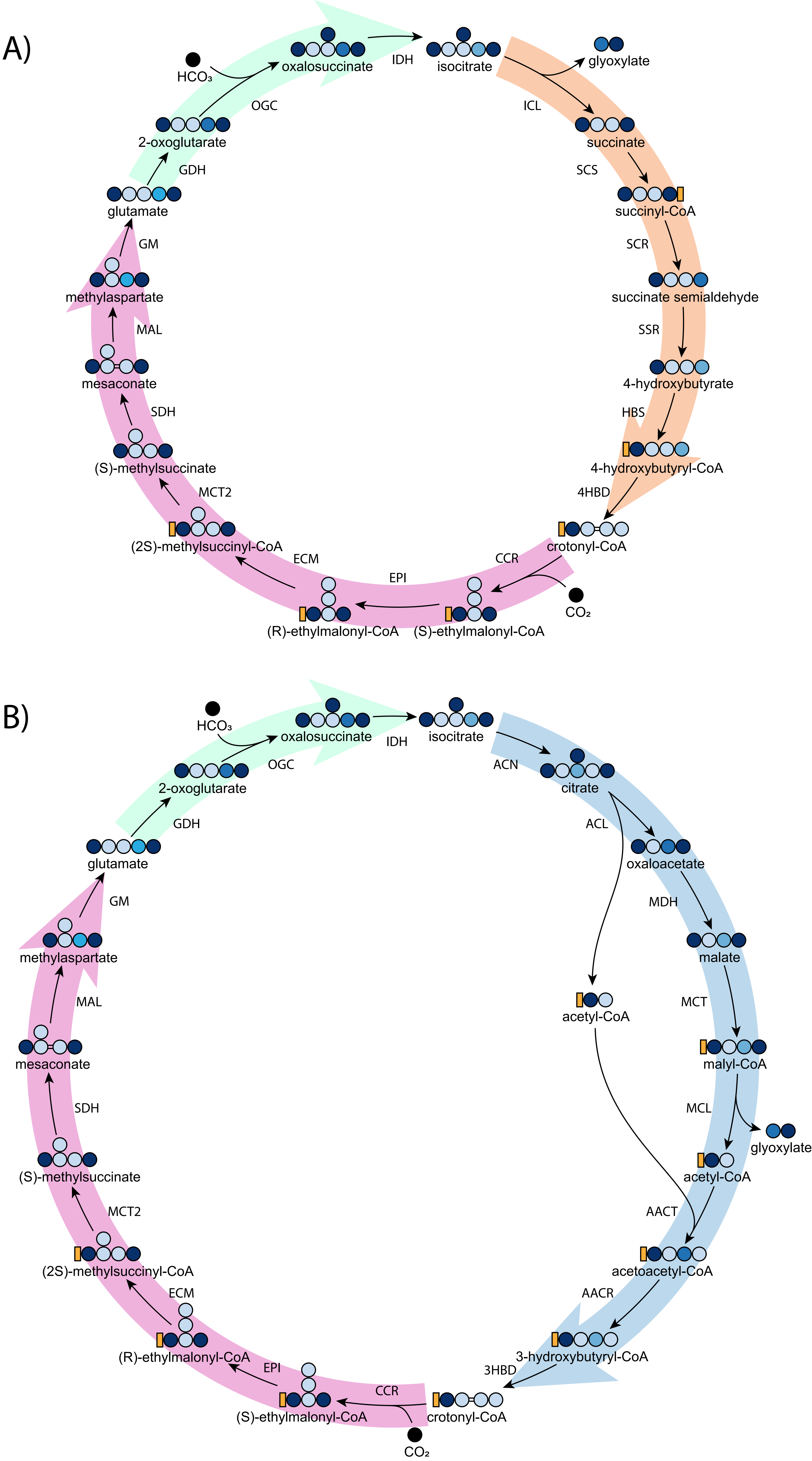

### Figure_2.pdf

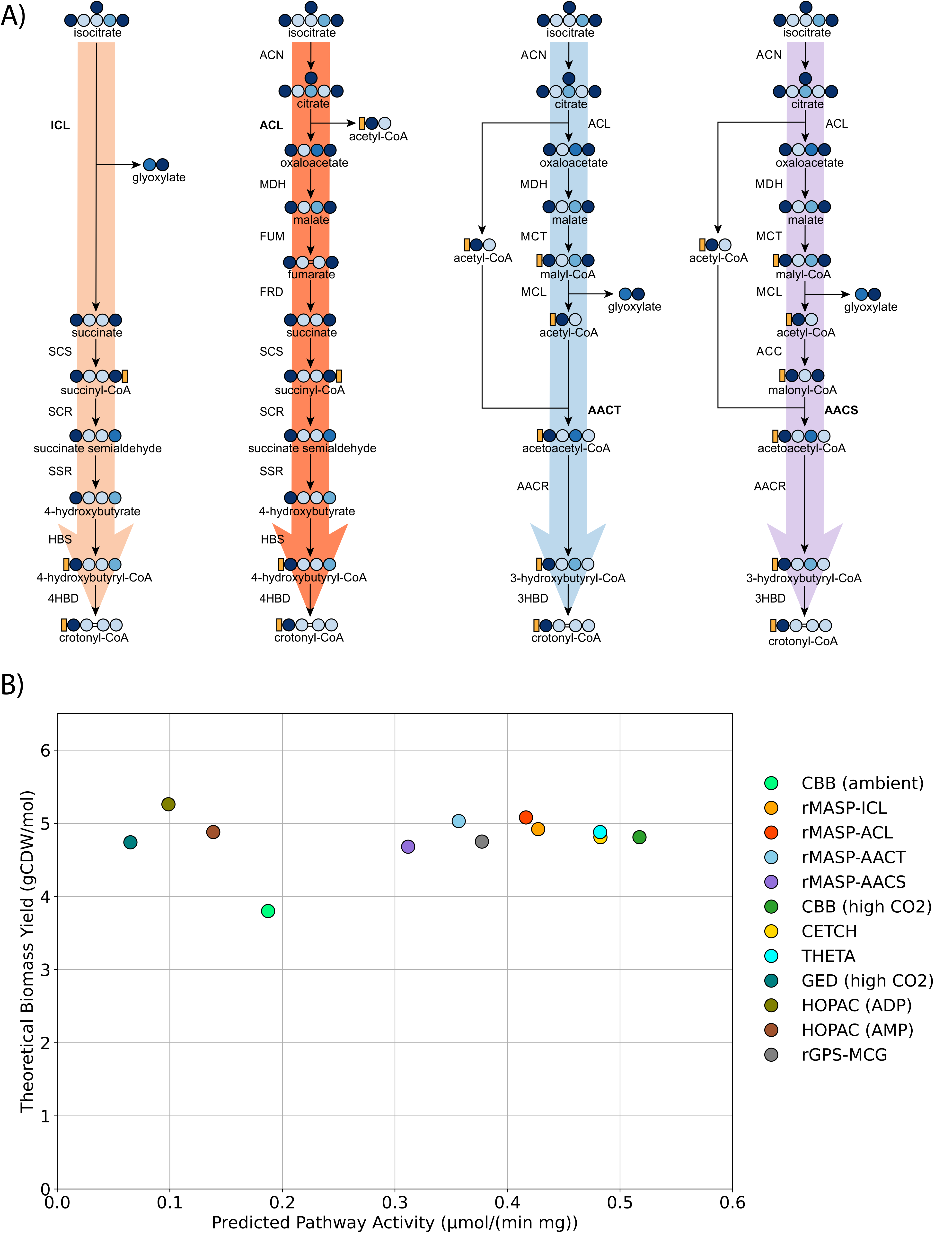

### Figure_3.pdf

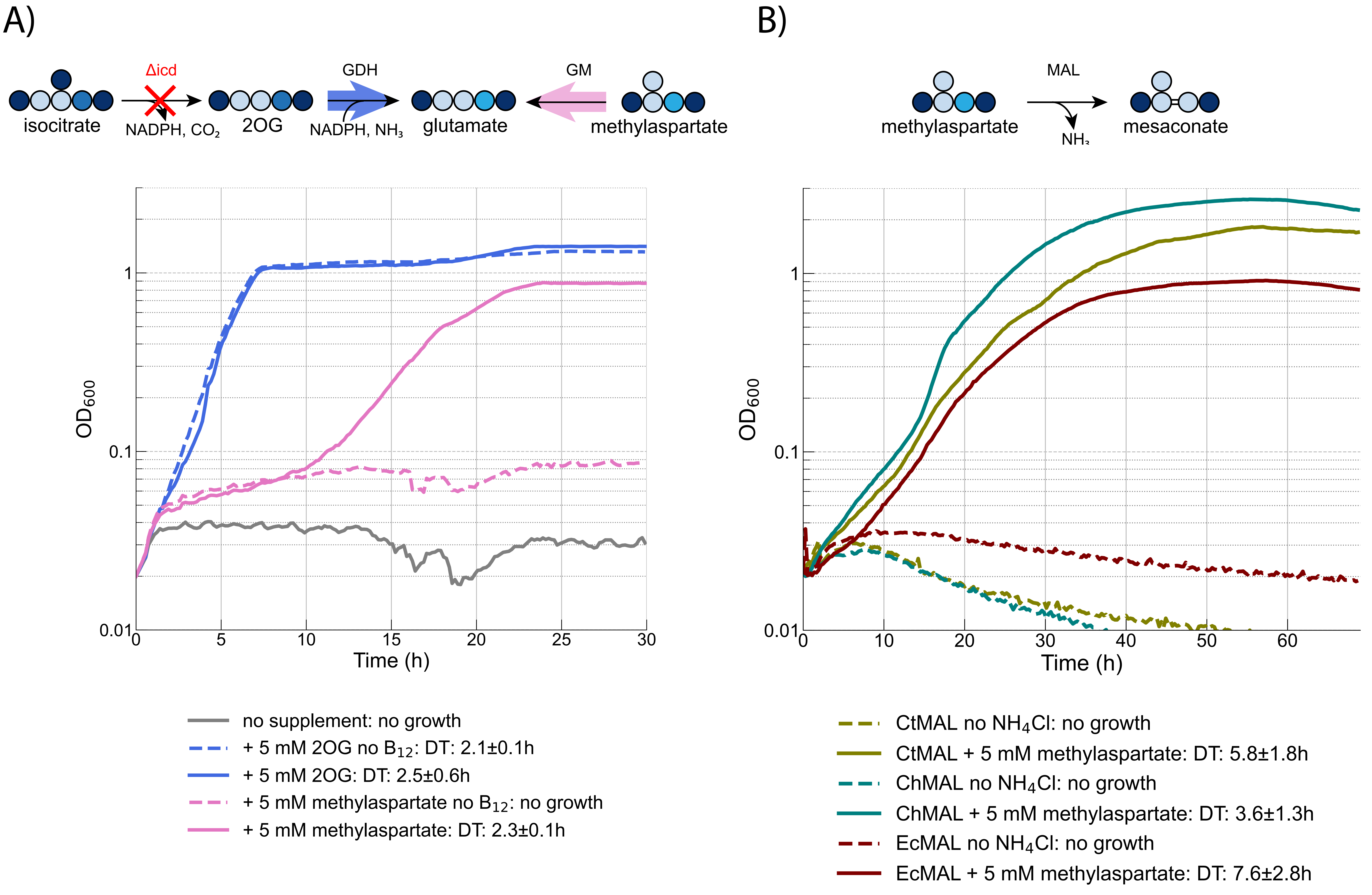

### Figure_4.pdf

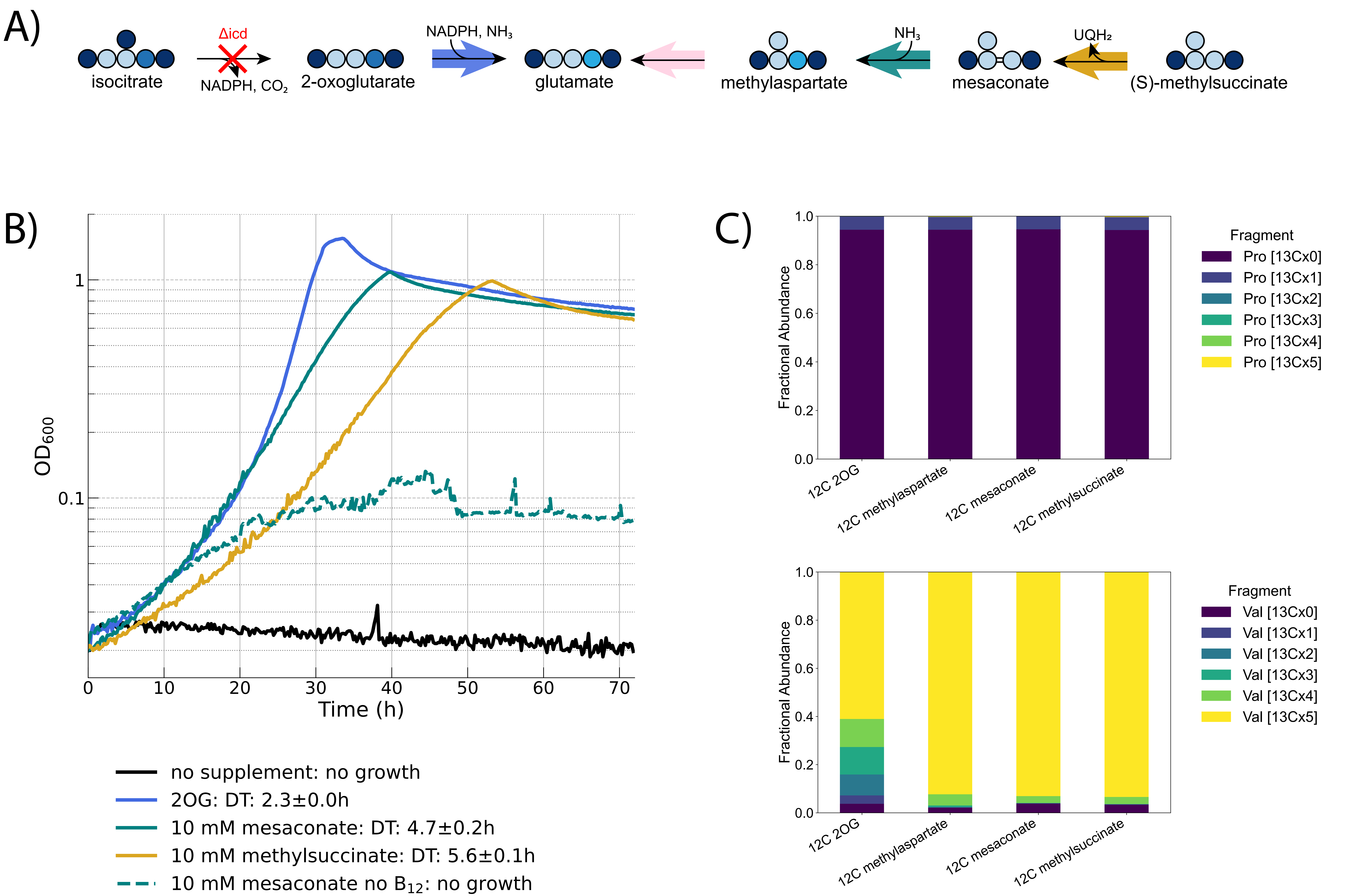

### Figure_5.pdf

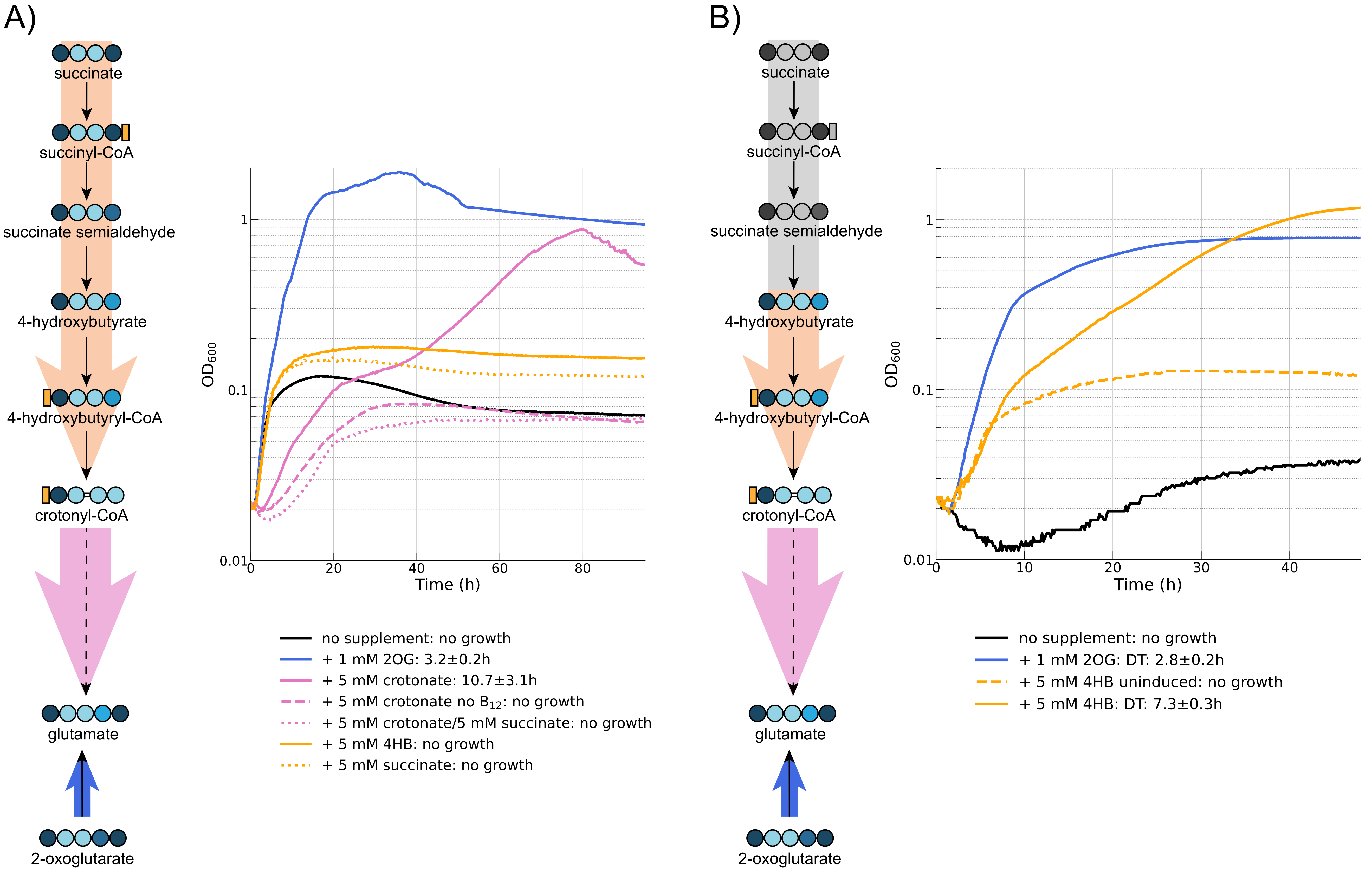

### Figure_6.pdf

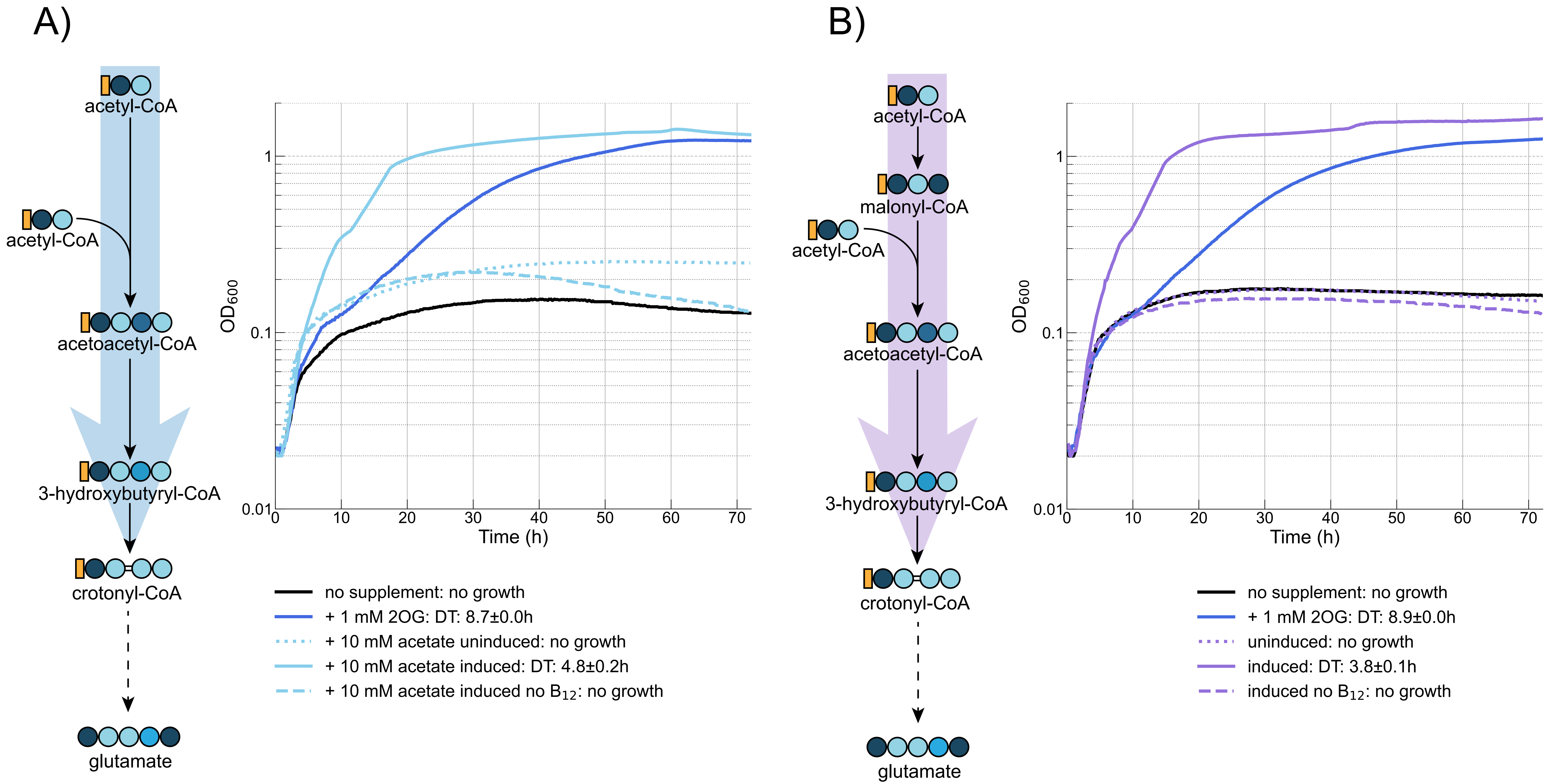

### Figure_7.pdf

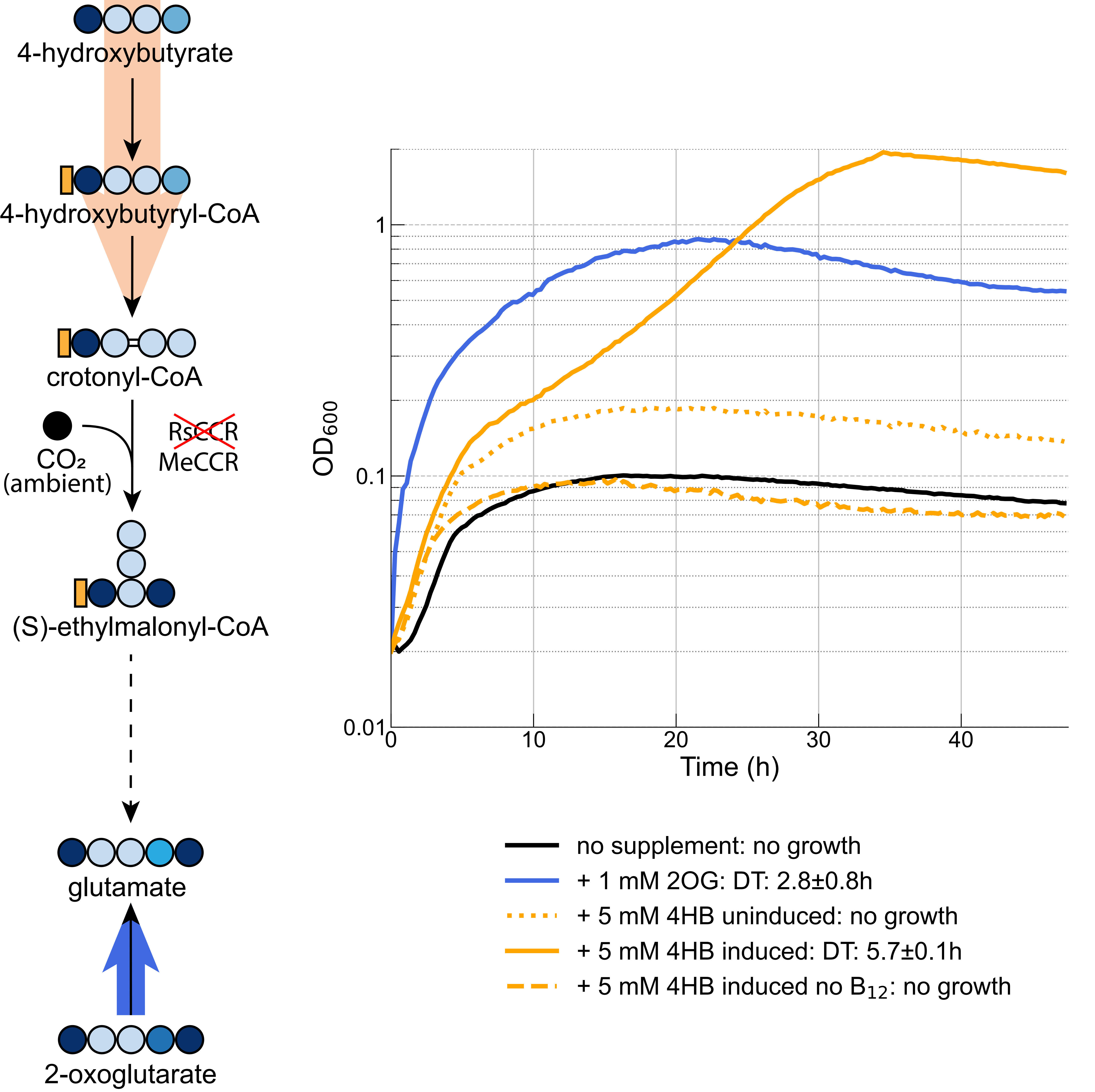

### Figure_8.pdf

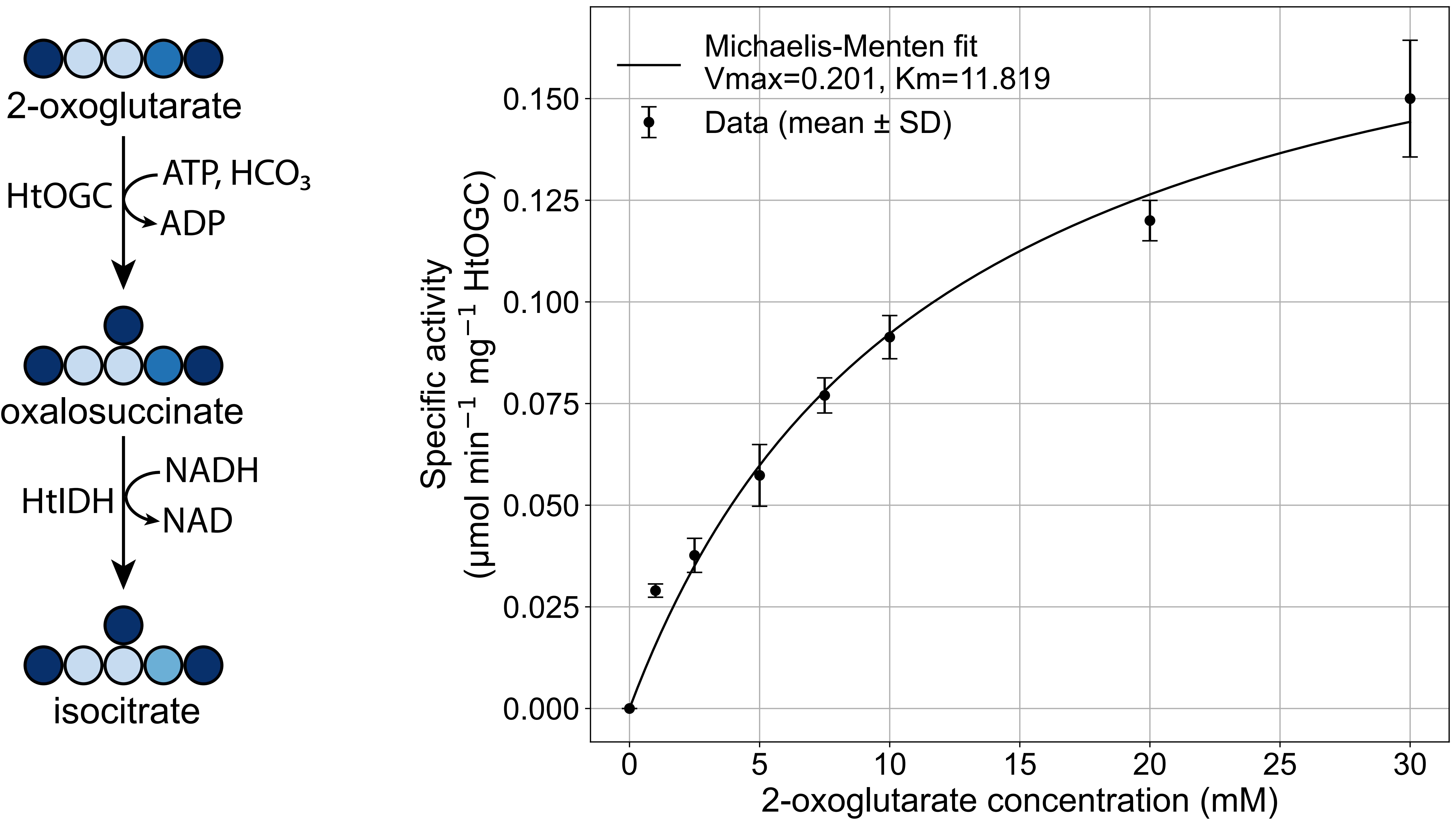

### Figure_S1.pdf

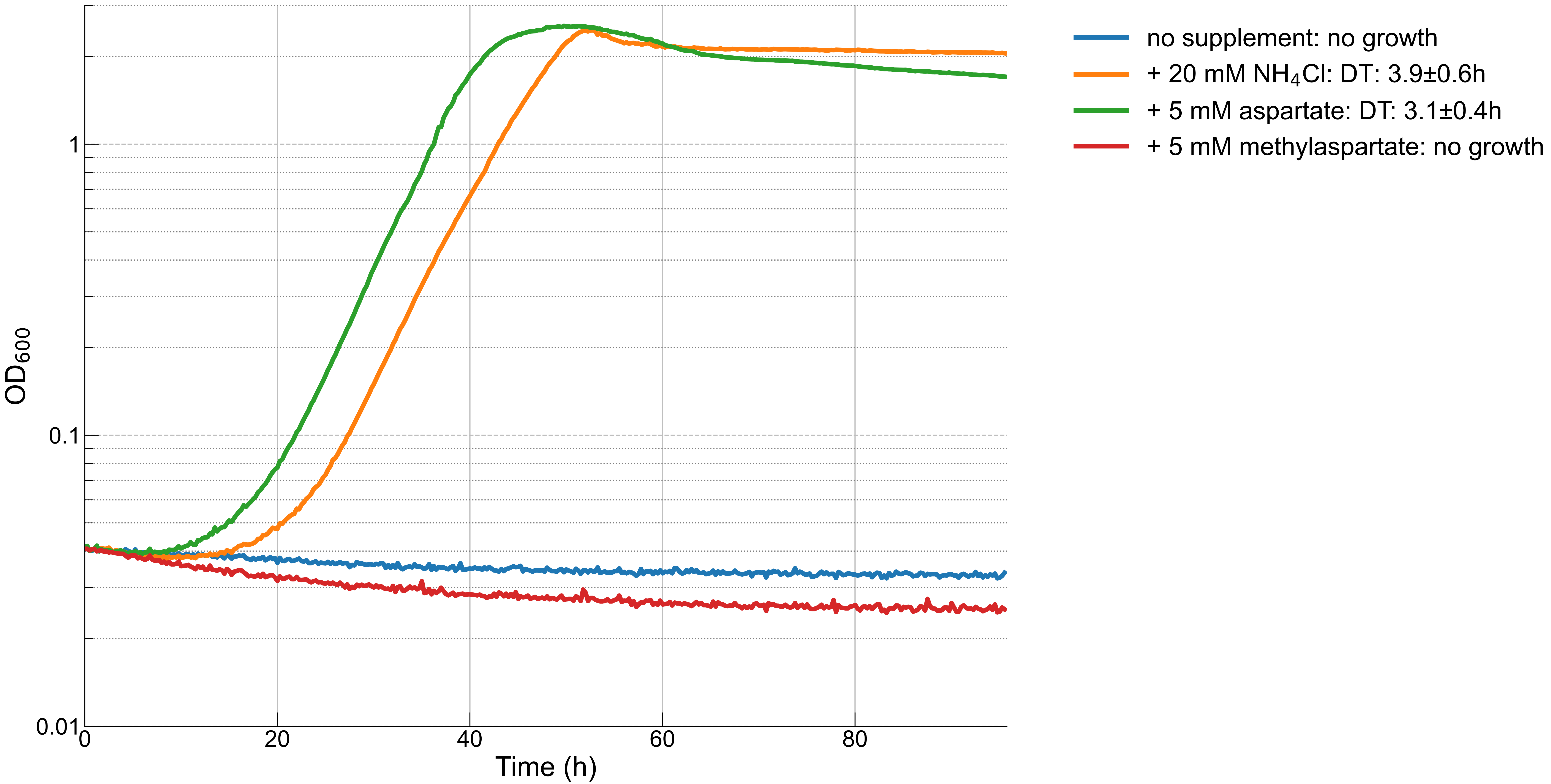

### Figure_S2.png

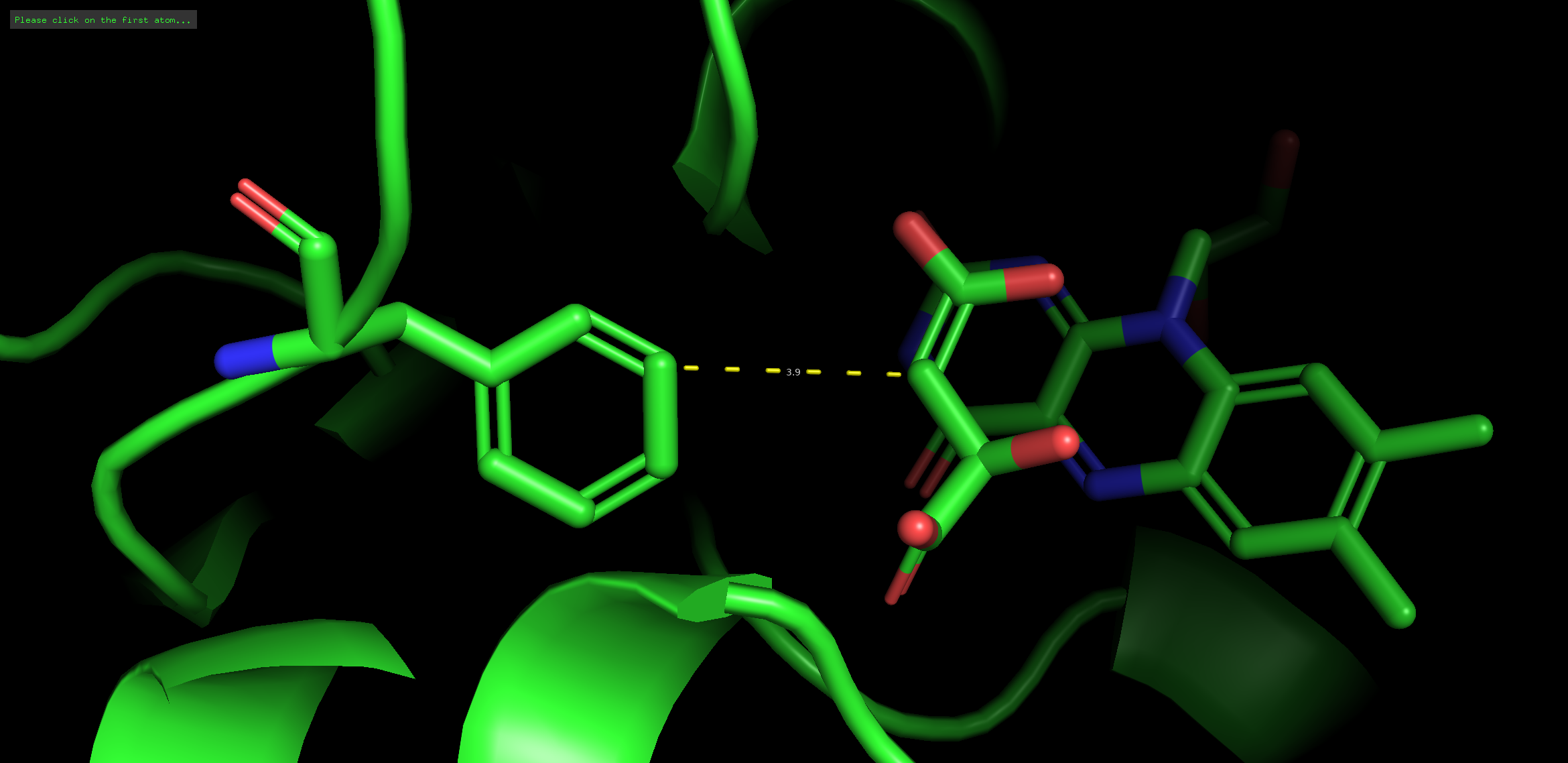

### Figure_S3.pdf

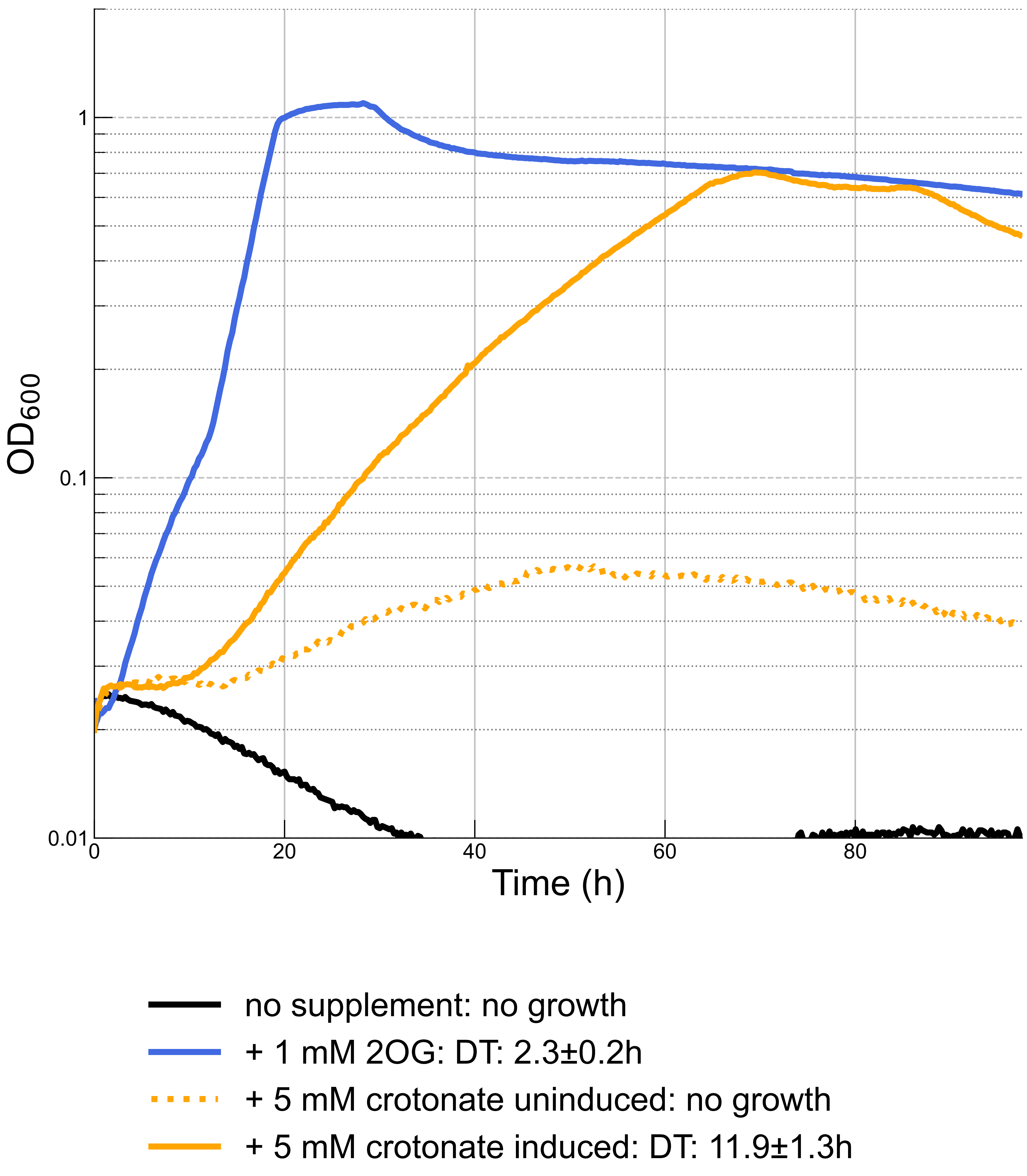

### Figure_S4.pdf

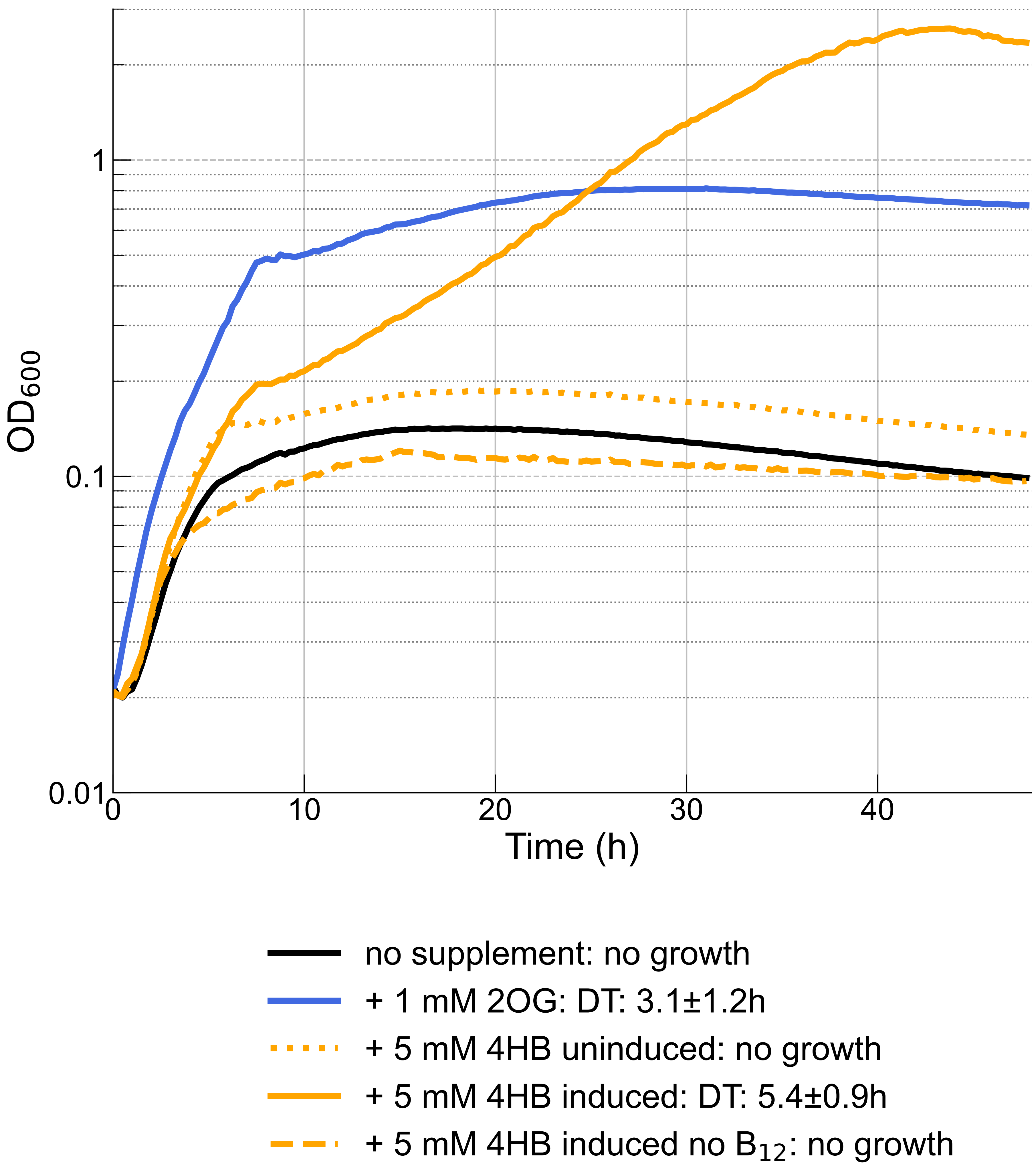

### Figure_S5.pdf

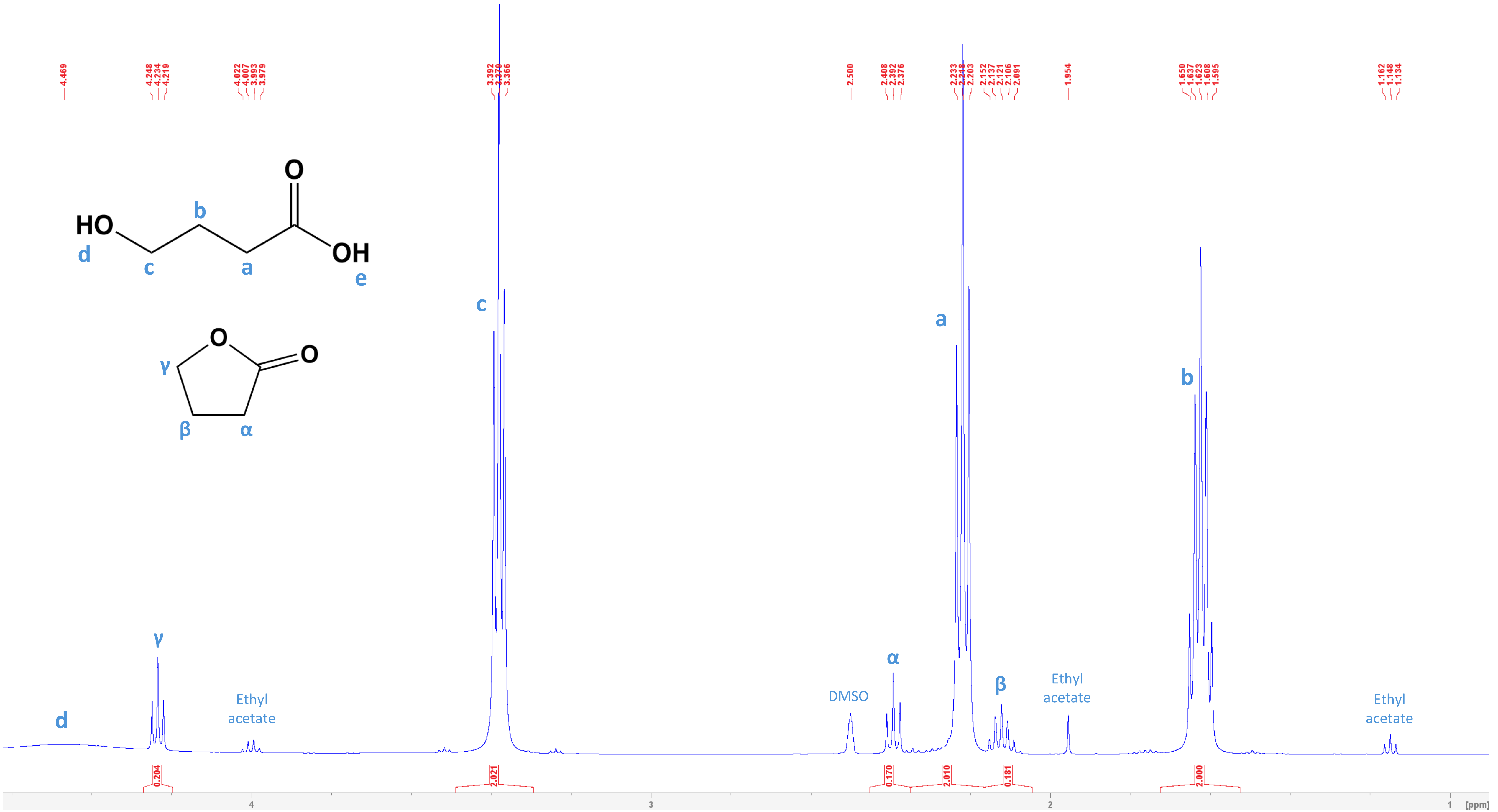

### Figure_S5.png

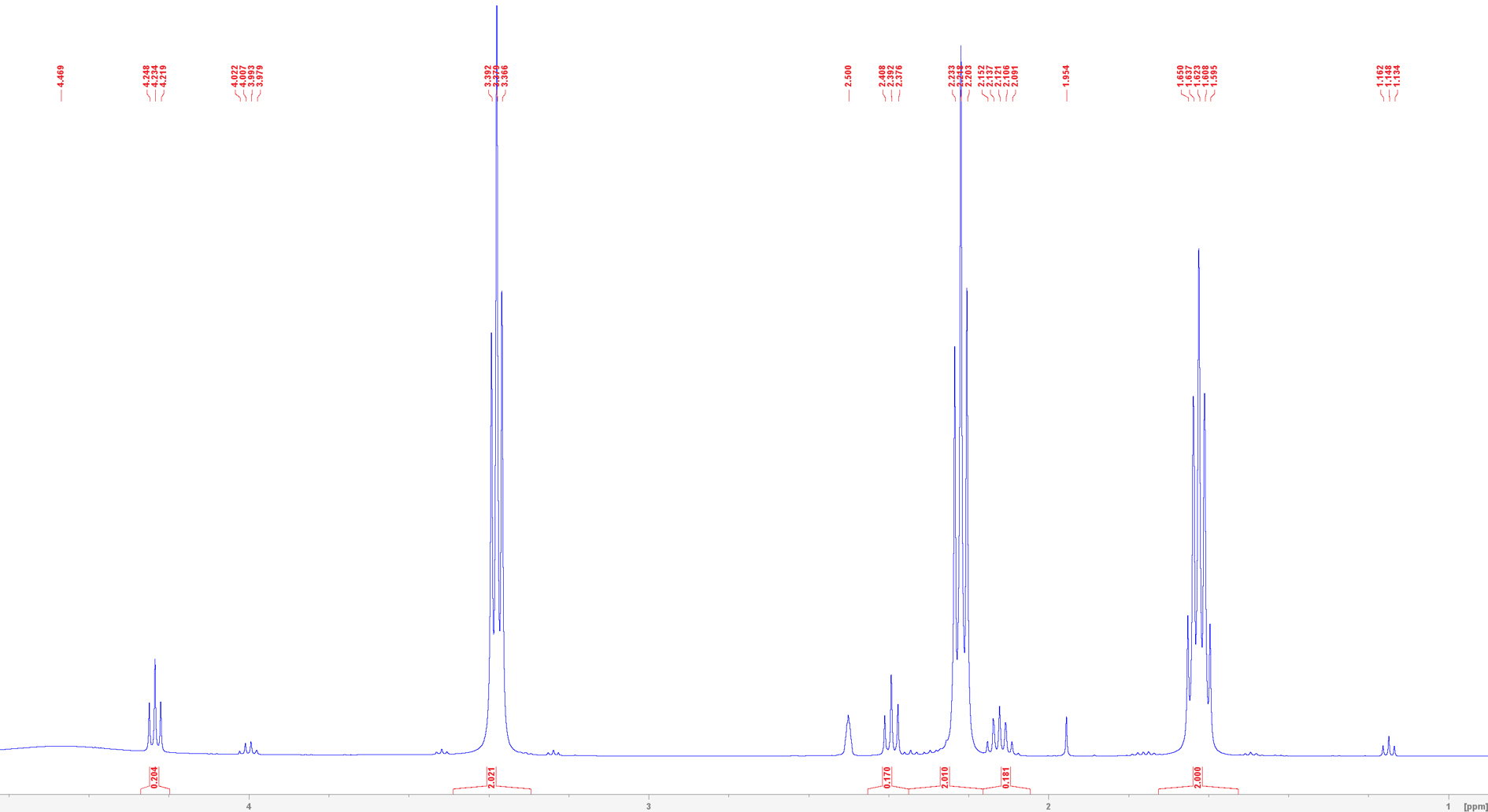
